# Structure and function of the bacterial and fungal gut flora of Neotropical butterflies

**DOI:** 10.1101/128884

**Authors:** Alison Ravenscraft, Michelle Berry, Tobin Hammer, Kabir Peay, Carol Boggs

## Abstract

The relationship between animals and their gut flora is simultaneously one of the most common and most complex symbioses on Earth. Despite its ubiquity, our understanding of this invisible but often critical relationship is still in its infancy. We employed adult Neotropical butterflies as a study system to ask three questions: First, how does gut microbial community composition vary across host individuals, species and dietary guilds? Second, how do gut flora compare to food microbial communities? Finally, are gut flora functionally adapted to the chemical makeup of host foods? To answer these questions we captured nearly 300 Costa Rican butterflies representing over 50 species, six families and two feeding guilds: frugivores and nectivores. We characterized the bacteria and fungi in guts, wild fruits and wild nectars via amplicon sequencing and assessed the catabolic abilities of the gut flora via culture-based assays.

Gut communities were distinct from food communities, suggesting that the gut environment acts as a strong filter on potential colonists. Nevertheless, gut flora varied widely among individuals and species. On average, a pair of butterflies shared 21% of their bacterial species and 6% of their fungi. Host species explained 25-30% of total variation in microbial communities while host diet explained 4%. However, diet was still relevant at the individual microbe level—half of the most abundant microbial species differed in abundance between frugivores and nectivores. Diet was also related to the functional profile of gut flora: compared to frugivores, nectivores’ gut flora exhibited increased catabolism of sugars and sugar alcohols and decreased catabolism of amino acids, carboxylic acids and dicarboxylic acids. Since fermented juice contains more amino acids and less sugar than nectar, it appears that host diet filters the gut flora by favoring microbes that digest compounds abundant in foods.

By quantifying the degree to which gut communities vary among host individuals, species and dietary guilds and evaluating how gut microbial composition and catabolic potential are related to host diet, this study deepens our understanding of the structure and function of one of the most complex and ubiquitous symbioses in the animal kingdom.

## Introduction

Microbes have been detected in the gut of almost every animal studied to date. This ubiquity is underpinned by the myriad functions these microbes serve: Gut microbes can assist animals with the uptake, synthesis and recycling of nutrients, breakdown of toxic or recalcitrant chemicals, and resistance to pathogens (Dillon and Dillon 2004). Despite their prevalence and importance, our knowledge of how and why these symbiotic communities change across host individuals, species, functional ecological groups, and geographical locations is still in its infancy.

At the most basic level, variation in the community composition of the gut flora can result either from exposure of the host to different pools of potential microbial colonists (including microbes transmitted vertically from parents to offspring), or from selective filtering of microbes by the physical and chemical conditions of the gut. Differences in diet or gut physiology between host dietary guilds, host species, and individual host organisms can affect both colonization by, and survival of, microbes in the gut.

Across disparate animal groups, host dietary guild has frequently attracted interest as a potential determinant of gut community composition. Different foods contain different microbial flora, and it is often assumed that consumption of foods can directly alter the composition of the gut community by introducing microbial colonists to the gut. This role appears to be most important during initial colonization. Once established the gut community is normally resilient, and most food-borne microbes pass through without becoming residents (e.g. Robinson et al 2010b; McNulty et al 2011). In addition to exposing hosts to alternative pools of microbes, diet can alter the composition of both horizontally acquired and vertically transmitted gut microbes by determining nutrient availability in, or affecting the chemical conditions of, the gut habitat (Robinson et al 2010a). Ingestion of a particular nutrient will promote the growth of gut microbes that digest that nutrient, leading to feedbacks between diet and microbial community function. For example, anaerobic fungi use lignocellulose as a food source; consumption of high fiber foods by ruminant species leads to greater abundance of these fungi than in non-ruminants (Gordon and Phillips 1998; Solomon et al 2016). These two mechanisms—food-borne microbial colonists and the chemical composition of the diet—are generally expected to result in similar gut flora among hosts that eat similar foods. Indeed, in many systems, gut community composition has been shown to be more similar between species that eat similar diets, even when these species belong to evolutionarily divergent groups (Colman et al 2012; Delsuc et al 2014). For example, herbivorous, omnivorous, and carnivorous mammals and fish host gut communities that are more similar within than between these three diet categories (Ley et al 2008a; Muegge et al 2011; Sullam et al 2012). In insects, species belonging to the wood-feeding and detritivorous guilds both host convergent gut flora (Colman et al 2012).

But host dietary guild is only one of many factors that may affect gut microbial community composition. Differences in colonization can also result from living in different habitats (e.g. Xiang et al 2006; Belda et al 2011), which results in sampling of divergent microbial pools by hosts. Additionally, the chemical conditions of the gut vary among species, independent of diet, and may act as a filter that selects for a specific gut community (Rawls et al 2006). For example, the guts of small animals, such as insects, range from fully aerobic to anaerobic and therefore favor different sets of microbes (Johnson and Barbehenn 2000). Similarly, gut pH varies among species and likely determines which microbes can establish in the gut (Beasley et al 2015).The host immune system can also regulate potential gut colonists (McFall-Ngai 2007; Salzman et al 2009).

Gut flora vary among host species not only in the composition of the core community, but also in the degree of variation among individuals. Such intraspecies variation can result from inter-individual differences in all of the factors mentioned above, including individual hosts’ diets, microbial source pools, gut chemistries, and immune systems. Species with the least variable communities are often those that depend on their gut flora to survive on particularly poor or recalcitrant diets, such as termites or herbivorous ants; these species also often have elaborately structured guts that both facilitate digestion and provide microenvironments that support a stable gut community (Engel and Moran 2013). In contrast, omnivorous hosts with simply-structured, tube-like guts, such as *Drosophila*, tend to have gut flora that vary more among individuals (Engel and Moran 2013). In general, gut membership is expected to be more stable when the gut flora serve a more crucial function for their host.

Insects offer a tractable system to understand the causes and consequences of variation in gut microbial communities due to their variation in dietary guild, their abundance and species diversity, and their often relatively simple gut flora. Further, insects are major primary consumers, pollinators, and disease vectors in terrestrial ecosystems. Their gut flora may therefore have a large, yet hidden, impact on ecosystem function (e.g. Nardi et al 2002). Indeed, the ability of many insects to subsist on nutritionally unbalanced or recalcitrant foods often stems directly from contributions of their gut flora (e.g. Warnecke et al 2007). Despite the importance of, and increasing interest in, the insect gut flora in general, the adult lepidopteran microbiome has been largely ignored. Although a few culture-based studies confirmed the presence of bacteria in the adult gut decades ago (Steinhaus 1941; Kingsley 1972), culture-independent profiling is only beginning. Such data exist for only a single species, *Heliconius erato* (Nymphalidae), and indicate that about 12 bacterial species dominate the adult gut (Hammer et al 2014).

In contrast to adults, the gut flora of several larval lepidopterans—particularly those of economic importance—have been characterized (Broderick et al 2004; Robinson et al 2010b; Pinto-Tomás and Sittenfeld 2011). However, the adult gut flora differ from those of larvae (Hammer et al 2014). This difference may derive from the fact that during pupation, the contents of the guts are voided, antimicrobial peptides are secreted into the gut lumen, and gut itself is replaced, presumably eliminating a large portion of the larval gut flora (Russell and Dunn 1996; Hakim et al 2010; Johnston and Rolff 2015). Indeed, in a culture-based study, Kingsley (1972) found that the density of colony-forming units derived from the guts of monarch butterflies (*Danaus plexippus*) decreased 1000-fold between pupae and freshly emerged adults. Existing data on larval lepidopterans therefore do not shed light on the gut flora of adult butterflies. Butterflies are biologically and economically important as herbivores and pollinators, and in areas outside of gut community ecology they are well-studied model organisms in the fields of population and nutritional ecology (Boggs et al 2003). Characterization of the causes and consequences of variation in the adult lepidopteran gut flora will therefore pave the way for connections between microbiome research and classical questions in ecology and evolutionary biology.

Here we use adult butterflies as a novel study system to characterize patterns of variation in the gut flora at the level of host individuals, species, and feeding guilds. We focus on how host nutritional ecology affects the community composition and functional capacity of the gut flora. The Neotropical community of butterflies allows for a high degree of replication both within and between species while controlling for geographic origin. Neotropical butterflies also exhibit well-described ecological variation that likely structures the gut community, particularly in feeding behavior: adults of some species feed on nectar, while others feed on the juice of rotting fruits. Adults are easily captured in the wild, permitting examination of the natural gut flora and avoiding the microbial community shifts frequently observed in captive animals (e.g. Chandler et al 2011). These characteristics make Neotropical butterflies a useful system to elucidation patterns in the composition and function of the gut microbial community.

We asked: (1) What bacteria and fungi are present in the guts of adult butterflies? (2) How does gut microbial community composition vary among individuals, species, and dietary guilds? (3) How does gut microbial composition compare to that of butterfly foods? (4) How does community composition translate into functional abilities—specifically, catabolism of various carbohydrates and amino acids?

Based on previous studies of other animal taxa (Ley et al 2008a; Colman et al 2012), we expected butterfly gut communities to cluster according to both host diet and host species, such that microbial beta diversity would be lowest between conspecific hosts, intermediate between hosts that belong to different species but the same feeding guild, and highest between heterospecific hosts that belong to different feeding guilds. We hypothesized that we would observe signals of environmental acquisition of the gut flora. Specifically, we predicted that most of the gut community would be a subset of the microbes present in the host’s food, and that frugivores, which feed on more “contaminated” foods (e.g. rotting fruit), would have more dense gut communities than nectivores. Finally, we hypothesized that the digestive abilities of the gut community would differ between feeding guilds, and that gut community catabolism should be highest for nutrients abundant in the host’s diet.

## Methods

### Study site

Butterflies were collected at La Selva Biological Research Station (10° 26’N, 83° 59’ W). La Selva is located in the lowlands on the Caribbean versant of the Cordillera Central in Costa Rica. The station owns 1,600 hectares, 55% of which is primary tropical wet forest; the remainder comprises secondary forest and disturbed habitats (e.g. abandoned pastures and plantations) in various states of regrowth. This habitat diversity allowed us to collect a large diversity of butterfly species while controlling for the geographic origin of our samples. La Selva also provided the facilities necessary for both the culture-dependent and -independent components of our project (e.g. a wet lab and laminar flow hood). From January to March 2013, we captured butterflies and sampled butterfly foods along the trail system in primary forest, secondary forest and recovering pastures. We returned to La Selva in March 2014 to collect additional food samples (fruits and nectars).

### Study organisms

Neotropical lepidopterans belong to two main feeding guilds: nectar feeders and fruit feeders. The Papilionidae, Pieridae, Lycaenidae, Riodinidae, Hesperiidae, and some Nymphalidae feed primarily on flower nectar, while several subfamilies of the Nymphalidae—Satyrinae, Morphinae, Charaxinae, and some members of the Nymphalinae—feed primarily on rotting fruits or other non-floral liquids (DeVries 1987, DeVries 1988). Our sampling included representatives of all of the nectivorous families—though most individuals derived from the Pieridae and Nymphalidae—as well as representatives from all the frugivorous nymphalid subfamilies described above (Supplementary Table 1). Three *Heliconius* (Nymphalidae: Heliconiinae) species were sampled; this genus supplements its nectivorous diet with pollen.

**Table 1.** Sample sizes. Number of butterfly or food samples (“individuals”) and species remaining in the sequencing datasets after rarefying.

|  | **Bacterial dataset** |  | **Fungal dataset** |  |
| --- | --- | --- | --- | --- |
|  | **Individuals** | **Species** | **Individuals** | **Species** |
| **Frugivores** | 148 | 24 | 92 | 15 |
| **Nectivores** | 142 | 28 | 69 | 19 |
| **Fruits** | 12 | 1 | 13 | 1 |
| **Nectars** | 46 | 8 | 39 | 8 |
| **Trap baits** | 25 | n/a | 14 | n/a |

### Sample collection and processing

We used aerial nets to catch both nectivorous and frugivorous species. To capture frugivores we also used Van Someren-Rydon traps (Daily and Ehrlich 1995) baited with a fermented mixture of fruit (primarily mango), molasses and rum. Traps were checked at least once every two days.

Upon capture, butterflies were placed in glassine envelopes and transported to the lab. We identified butterflies to species using *The Butterflies of Costa Rica and Their Natural History* (Devries 1987). Butterflies were fed a 5 to 10 microliter droplet of a filter-sterilized solution of sugar and nigrosin dye (Sigma 198285) to assist with visualization of their guts during dissection. Butterflies were euthanized with ethyl acetate, washed with 70% ethanol, and dissected in a laminar flow hood to minimize contamination with environmental bacteria and fungi. Guts were removed and homogenized in a small amount of inoculating fluid (described below) in a sterile 2 mL tube. A portion of the homogenate was cultured to assess gut community catabolic profile (described below). The remainder of the gut homogenate was suspended in cetyl trimethyl ammonium bromide (CTAB) and refrigerated at 4°C until transportation to Stanford University for DNA extraction. CTAB is an effective preservative for storage of insect samples prior to microbiota analysis (Hammer et al 2015).

We also collected samples of the butterflies’ foods to characterize potential source pools of microbial gut communities. Trap baits were sampled in 2013 and 2014 by suspending approximately 0.25 to 0.5 mL of rotten fruit slurry in CTAB. In 2014, nectar samples were also collected. We used capillary tubes to extract nectar from flowers in the field, obtaining between 0.5 and 10 μL per flower. Nectars were also preserved in CTAB. We only collected nectar from species at which we directly observed at least one instance of butterfly feeding. In 2013, we sampled fruits of *Dipteryx oleifera* (Fabaceae) by scraping flesh from the surface of the fruit into CTAB. In 2014, we rinsed the fruit with 5 mL of sterile water and preserved 0.25 to 0.5 mL of the rinsate, since butterflies only feed on surface juices and do not consume the pulp. *Dipteryx oleifera* is the second-most abundant leguminous tree in primary forest at La Selva (McDade et al 1994) and was the only wild fruit we observed butterflies feeding upon during both visits to the site.

Research was carried out under permit numbers 202-2012-SINAC and R-016-2014-OT-CONAGEBIO.

### Community catabolic profiling

A portion of each gut homogenate was seeded into 10 mL of inoculating fluid (IF-A, Biolog Catalog No. 72401) and cultured in microbial catabolic phenotype plates (Gen III microplates, Biolog Inc., Hayward, CA). The volume of gut homogenate used depended on the size of the butterfly: we used 25 microliters for most animals, but took less volume from the largest species (whose gut homogenate was more concentrated, since the guts themselves were large), and more volume from the smallest species (whose gut homogenate was less concentrated, since the guts were small). This was meant to correct for the grossest differences in inoculation density; further corrections are described below.

The catabolic phenotyping plates allowed simultaneous assessment of catabolism of 71 carbon and nitrogen sources (e.g. glucose, fructose, ammonia, uric acid; Table S5) via a colorimetric assay. Plates were incubated at ambient temperature. Absorbance at 590 nm was measured twice per day using a 96-well plate reader (Chromate-4300, Awareness Technology Inc., Palm City, FL) until color remained stable (5 days on average).

To reduce bias due to differences in inoculum density between the samples, the data were standardized as in Garland et al (2001). Briefly, based on visual inspection of the distribution of plate averages, we chose a target average absorbance value per plate of 190. For each plate, we selected data from the timepoint at which the average absorbance was closest to this target. These measurements were standardized by subtracting the value of the negative control and dividing by the plate’s average absorbance.

### Microbial community characterization

Samples were shipped to Stanford University, where they were homogenized via bead beating, extracted with chloroform, and cleaned using the DNeasy Blood and Tissue Kit (Qiagen, Germantown, MD). Our DNA extraction protocol is described in detail in Peay et al (2007).

Bacterial DNA was amplified with primer set 515f (5’-GTGCCAGCMGCCGCGGTAA-3’) and 806r (5’-GGACTACHVGGGTWTCTAAT-3’), which amplifies the V4 hypervariable region of the 16S rRNA with few taxonomic biases (Bergmann et al 2011). The bacterial amplicons were indexed with barcoded forward and reverse primers to (Caporaso et al 2012; Kozich et al 2013). The PCR reaction contained 0.2 μM forward primer, 0.2 μM reverse primer, 0.2 mM dNTP, 0.65 U OneTaq HotStart (New England Biolabs) and 1X Thermopol buffer (New England Biolabs) in a volume of 25 μL. The thermocycler program began with denaturation at 94 C for 3 minutes followed by 35 cycles of denaturation at 94 C for 45 seconds, annealing at 50 C for 60 seconds, and extension at 68 C for 90 seconds, with a final extension of 68 C for 10 minutes (adapted from the Earth Microbiome Project protocol; Gilbert et al 2014).

Fungal DNA was amplified with primers ITS1f (5’-CTTGGTCATTTAGAGGAAGTAA-3’) and ITS2 (5’-GCTGCGTTCTTCATCGATGC-3’) (White et al 1990; Gardes and Bruns 1993). The reverse primer was barcoded to index the samples (Smith and Peay 2014). The PCR recipe was identical to the bacterial PCR recipe above. Thermocycler conditions were denaturation at 95 C for 1 minute followed by 35 cycles of 94 C for 30 seconds, 52 C for 60 seconds, and 68 C for 60 seconds, with a final extension of 68 C for 5 minutes. Each bacterial and fungal amplification was run in triplicate to minimize the effects of stochastic amplification.

Bacterial and fungal PCR products were cleaned with AMPure XP magnetic beads (Agencourt A63881) and the final concentration of each sample was quantified fluorometrically with a 96-well plate reader (Qubit dsDNA HS kit, Thermo Fisher Q32854). An equal mass of DNA from each sample was added to the bacterial or fungal library, respectively. Bacterial and fungal libraries were sequenced in separate runs on an Illumina MiSeq platform at the Stanford Functional Genomics Facility (Stanford, CA), with 2 × 250 chemistry for bacteria and 2 × 300 chemistry for fungi.

### Illumina sequence data processing and cleaning

We used the program cutadapt (Martin 2011) to remove priming sites and poor quality bases at the 5’ and 3’ ends of the sequences. Sequences were merged and clustered at a 97% similarity cutoff with UPARSE (Edgar 2013). Both de-novo and reference-based chimera checking were performed in UPARSE; bacterial reads were compared to the RDP Gold database and fungal reads to the UNITE database. Bacterial taxonomy was initially assigned using the RDP classifier (Wang et al 2007) with Greengenes (McDonald et al 2012) as the training set. We used the RDP classifier with the Warcup (Deshpande et al 2016) training set to assign fungal taxonomy. We checked and revised these assignments by aligning our representative sets of bacterial and fungal sequences against the NCBI nucleotide collection using BLAST.

OTUs that were identified as lepidopteran 18S, archaeans, mitochondria, or chloroplasts were excluded. (These accounted for less than 0.1%, less than 0.1%, 0.6%, and 4.8% of the raw total reads, respectively, and 98% of the chloroplast reads derived from samples of butterflies’ foods.) Additionally, DNA extraction kits and other laboratory reagents are known to contain microbial DNA that can contaminate microbiome analyses (Salter et al 2014). We found fourteen bacterial OTUs that were present at higher abundance in the negative controls than the butterfly gut samples (Supplementary Table 3); these were classified as contaminants and removed from the dataset prior to statistical analyses. The fungal dataset (especially nectivore samples) contained many ectomycorrhizal lineages that were unlikely to be members of the butterfly gut community. This suggested that the butterfly gut samples (particularly those of nectivores) had low fungal biomass, resulting in detection of ambient lab and environmental contaminants. We removed lineages that were likely contaminants from the analysis: we omitted 135 OTUs assigned to the genus *Mortierella*, which are decay fungi that inhabit soil. Wood decomposers and ectomycorhizal, arbuscular or ericoid mycorrhizal fungi were also removed from the dataset, as were lineages that were present at higher abundance in the negative controls than in the samples (Supplementary Table 4). We retained lineages that were known phylloplane inhabitants, plant pathogens, and fruit decomposers, as well as most ascomycete and basidiomycete yeasts.

Frugivorous butterflies were captured either with traps baited with fermented fruit or with aerial nets. For those captured in baited traps, a portion of the gut flora likely derived from microbes in the trap baits. To identify these microbes, we tested for differential abundance of bacterial and fungal OTUs between trapped frugivores and netted frugivores using the *mt* function in the R package “phyloseq” (McMurdie and Holmes 2013), which includes an FDR correction for multiple testing. No bacterial OTUs differed in abundance between netted and trapped animals, but three fungal OTUs were significantly more abundant in the trapped butterflies. These OTUs were all strains of the yeast *Kazachstania exigua*, which was also by far the most abundant fungus detected in the traps. These three OTUs accounted for 43.7% of all fungal reads from the fermented baits and only 0.3% of reads from wild fruits. We concluded that the OTUs had been introduced into the trapped butterflies’ guts via the fermented baits and removed them from the data derived from butterfly samples.

To control for differences among samples in sequencing depth, we rarefied the sequencing data using phyloseq. Bacteria were rarefied to 1000 and fungi to 200 sequences per sample. These cutoffs allowed us to retain as many samples in the dataset as possible while still profiling the dominant microbes. All sequencing results reported are based on rarefied data, unless otherwise noted.

### Quantification of total bacterial abundance

Bacterial DNA was quantified with qPCR using SYBR green fluorescent chemistry (iCycler IQ, Bio-Rad, Hercules, CA). Since the 515f/806r primer set sometimes amplifies butterfly 18S in addition to bacterial 16S rRNA, we designed a PNA clamp to block 18S amplification (Lundberg et al 2013). The clamp sequence was GCCCGCTTTGAGCACTCT and it was synthesized by PNA Bio (Thousand Oaks, CA). The reaction volume was 20 μL and the recipe was 7.5 μM PNA clamp, 0.2 μM 515f primer, 0.2 μM 806r primer, and 1X PerfeCTa SYBR Green FastMix for iQ (Quanta Biosciences, Gaithersburg, MD). Thermocycler settings were an initial 10-minute denaturation at 95 C followed by 45 cycles of denaturation at 95 C for 15 sec, PNA clamp annealing at 76 C for 10 sec, primer annealing at 50 C for 30 sec, and extension at 68 C for 30 sec. To check that amplified fragments were the expected length, we performed a melt curve ramping from 55 C to 95 C in 0.5 C increments at 10 second intervals. PCR products were also visualized using gel electrophoresis.

To calculate the starting number of 16S rRNA copies, each sample’s threshold cycle was compared to an internal standard curve ranging from an initial 10 to 10^7^ copies per μL of *E. coli* 16S rRNA amplicons. (Generation of these standards is described below.) Each sample and standard was run in triplicate and the results were averaged. The correlation coefficients of the standard curves for all qPCR plates ranged from 0.973 to 0.988.

Since a fraction of each sample was removed for culture-based catabolic profiling prior to DNA preservation and extraction, the qPCR estimates were corrected accordingly. For example, if 50% of the gut homogenate from a given sample was removed prior to DNA preservation, that sample’s qPCR estimate was multiplied by two to obtain the estimated total 16S rRNA count for the entire gut.

The 16S rRNA standard curve was generated as follows: 16S rRNA was amplified from *E. coli* using primers 27f and 1492r, ligated into a plasmid vector, and cloned into chemically competent cells. Colonies were screened for inserts of the expected size by PCR amplification using M13f and M13r primers followed by gel electrophoresis. To further screen for the correct insert, several colonies were bidirectionally sequenced. One colony that passed both screens was selected for the standard curve. This clone was grown to saturation in LB + kanamycin selective media. Plasmids were extracted using the Qiagen Plasmid Mini kit and linearized using the restriction enzyme Spel (FastDigest, Thermo Fisher FD1253). DNA was purified using a Qiagen PCR cleanup column and run on a gel to verify complete linearization. DNA concentration was quantified with PicoGreen and copy number per μL was calculated as the molecular weight of plasmid plus insert (g/molecule, calculated as length in bp * 660 Da/bp / 6.02*10^23^) divided by the measured DNA concentration (g/μL). Via serial dilution of the raw extract with sterile TE, we generated a standard curve ranging from 10^7^ copies/μL to 10 copies/μL.

We were not able to quantify total fungal abundance because our ITS primers amplified fragments of vastly different lengths (ranging from 250 base pairs to over 1000 bp).

### Statistical analyses

All models had Gaussian error structure and were fit in R version 3.2.2 (R Core Team 2015). Models without random effects were fit using the *lm* command in the “stats” package. Models with random effects were fit using the *lmer* command in the package “lme4” (Bates et al 2014).

#### a. Variation in total bacterial load among host species and feeding guilds

A log transformation was applied to the 16S rRNA counts prior to analysis. We used linear and linear mixed effect models (Zuur et al 2009) to assess whether total bacterial load differed among host species or between host feeding guilds. We used a likelihood ratio test to compare models with and without a random effect of host species, including fixed effects of host diet, wing length (as a proxy for the animal’s size), and capture date. We then tested for significance of the fixed effects using backwards model selection with likelihood ratio tests, including a random effect of host species in all models.

#### b. Variation in OTU richness

To test whether observed bacterial and fungal richness differed among host species, we again used a likelihood ratio test to compare models with and without a random effect of host species, including a fixed effect of host diet to control for overall diet-based differences. We tested for correlation between host diet and observed bacterial and fungal richness by comparing models with a fixed effect of diet to models without the fixed effect; both models had a random effect of host species. We omitted data from host species represented by a single individual from these analyses.

#### c. Variation in microbial community composition

##### Between feeding guilds

To test for a relationship between butterfly feeding guild and gut microbial community composition, we used ordination (NMDS) plots to visualize and perMANOVAs to test for dissimilarity in microbial community composition (Anderson 2001). Both operations were performed on the Bray-Curtis dissimilarities between rarefied samples. We performed two perMANOVAs (the *adonis* test in the R package “vegan”; Oksanen et al 2015) to test for dissimilarity in bacterial and fungal communities, respectively. These tests included terms for both host diet and host species, thus simultaneously testing whether gut communities differed between host species and feeding guilds. Host species represented by a single individual were omitted from these analyses.

If communities differ in variance, perMANOVA results can be unreliable (Anderson and Walsh 2013). To verify the results of the tests, we therefore tested for differences in dispersion among feeding guilds using the *betadisper* function from the package “vegan.” For bacteria, the feeding guilds did not differ in dispersion (anova: *df* =1, F=0.12, p=0.73) therefore the data conformed to the assumptions of perMANOVA. For fungi, the feeding guilds did differ in spread (anova: *df* =1, F=4.4, p=0.04). However, after removing the five host species with extreme dispersions, the feeding guilds no longer differed in spread (anova: *df* =1, F=1.3, p=0.21). The perMANOVA results with these five species removed were qualitatively identical and numerically similar to those obtained from all host species, therefore we report the test results for all species (omitting those represented by a single individual).

In order to understand the effect of host diet on individual OTU abundances, we modeled the relative abundance of a bacterial or fungal OTU (counts out of 1000 or 200, respectively) as a function of the interaction between host diet and OTU identity, with a random effect of host species. Since it was not feasible to model abundances of every OTU, we selected the 20 bacterial or 20 fungal OTUs that were present at the highest abundances in the pooled data and were detected in at least 10% of either frugivore or nectivore samples. Prior to the analyses, abundance data were transformed by log(x+1) to homogenize variance.

##### Among host species

We used perMANOVAs (described above) to test for dissimilarity in the gut flora among host species while accounting for differences between feeding guilds. Since perMANOVA assumes homogeneity of variance among communities, we again used the *betadisper* function to test for differences in dispersion. Host species’ bacterial and fungal communities did differ in dispersion (bacterial anova: *df* =36, F=2.4, p< 0.001; fungal anova: *df* =28, F=11.4, p< 0.001). For bacteria the difference was driven by five species that had unusually low variance; when these were removed, the remaining 32 species no longer differed in spread (anova: *df* =31, F=0.9, p=0.65). For fungi, the difference was also driven by five extreme species. The remaining 24 species in the fungal dataset no longer differed in variance (anova: *df* =23, F=1.3, p=0.16). For both bacteria and fungi, we re-ran the perMANOVA tests using the subset host species that did not differ in dispersion. Results were again qualitatively the same as those for all host species, so we report the results for all species.

##### Among individuals

To quantify beta diversity among individual butterflies’ gut flora, we calculated the median percentage of OTUs shared between individuals, where the percentage shared between individuals A and B was calculated as 0.5 * (number of A’s OTUs shared by B / total OTUs in A + number of B’s OTUs shared by A / total OTUs in B).

##### Between food microbial communities and gut communities

We used NMDS plots (on Bray-Curtis distances between rarefied samples) to visualize differences between the butterfly gut flora and the microbial composition of butterfly foods. To test whether gut flora were more similar to the microbial communities of one food than to another food, we calculated the pairwise Bray-Curtis distances between each butterfly gut community and each food community and performed t-tests on the per-butterfly averages of these distances for the three foods (fruits, nectars and baits). To correct the resulting p-values for multiple testing, we used the Benjamini-Hochberg false discovery rate (FDR) method as implemented by the *p.adjust* function in R (Benjamini and Hochberg 1995).

#### d. Variation in microbial community function

The Hellinger transformation was applied to the standardized catabolic data prior to all analyses.

##### With gut community composition

To test whether differences in microbial community function were correlated with differences in microbial species composition, Mantel tests were performed between the Bray-Curtis distances between bacterial or fungal microbial communities and the Euclidean distances between catabolic profiles. Since trap bait-derived fungal OTUs (three strains of *Kazachstania exigua*) were introduced into the Biolog plates via frugivores’ guts, these OTUs were included in the calculation of Bray-Curtis distances between fungal communities. However, results were qualitatively equivalent when these three OTUs were omitted from the Bray-Curtis calculations.

##### With host species and diet

First, we tested for correlation between gut community catabolic profile and both host diet and host species by performing a perMANOVA on the Euclidean distances between the gut catabolic profiles. (Singleton host species were omitted from this analysis.) To investigate differences between frugivores and nectivores in more detail, we next modeled substrate catabolism as a function of the interaction between host diet and substrate identity, including a fixed effect for the log of the number of 16S rRNA copies in the host’s gut to control for differences in bacterial density, and a random effect for the identity of the host butterfly to control for overall differences in catabolic activity of the gut community among individuals. Finally, to test whether catabolism of certain classes of substrate (for example, amino acids or sugars) was consistently up- or down-regulated with host diet, we modeled substrate catabolism as a function of the interaction between host diet and substrate class, with a fixed effect for log(16S rRNA counts) and a random intercepts of host identity as before, plus an additional random intercept and random diet effect (slope term) for each substrate. Thirteen substrates belonged to classes with three or fewer substrates and were therefore omitted from the class-level analysis. For both models, least-square means and the contrasts between them were calculated using the *lsm* command in the “lsmeans” package (Lenth 2016) to assess the whether a substrate or substrate class was catabolized differently between frugivores and nectivores. The resulting p-values for each substrate or substrate type, respectively, were FDR corrected to control for multiple testing.

## Results

Below we first describe the overall composition of the adult butterfly gut flora, then address patterns of variation in microbial community composition at the level of host feeding guild, host species, and host individual. Gut communities are then compared to the microbes present in butterfly foods. Functional differences between gut microbial communities are described last.

### 1. Bacteria and fungi present in the butterfly gut

We sequenced the bacterial flora of 306 butterflies, of which 290 were retained after rarefaction, and the fungal gut communities of 247 butterflies, 161 of which were retained after rarefaction. (See Table 1 for a full sample size breakdown and Supplementary Table 1 for counts per butterfly host species). After sequence processing, quality filtering, and removal of contaminant OTUs, 7.2 million bacterial sequences and 1.5 million fungal sequences were obtained from butterfly guts. After rarefying, a total of 958 bacterial OTUs and 880 fungal OTUs were observed across all butterflies. (Note that numbers of bacterial and fungal OTUs cannot be compared because bacteria and fungi were rarefied to different read depths.) Individual butterflies hosted a mean of 30 (interquartile range:18-37) bacterial OTUs (number observed at a 1000 read per sample cutoff) and 21 (IQR: 12-28) fungal OTUs (number observed at a 200 read per sample cutoff).

The most prevalent bacterial phyla and classes across all butterflies were Proteobacteria:Gammaproteobacteria (40% of all butterfly-derived bacterial sequences), Proteobacteria:Alphaproteobacteria (28%), Firmicutes:Bacilli (12%), Bacteroidetes:Flavobacteriia (7%), and Tenericutes:Mollicutes (4%). The dominant fungal class was Saccharomycetes (62% of all butterfly-derived fungal sequences) in the phylum Ascomycota. All of these Saccharomycetes belonged to the order Saccharomycetales, which defines the “budding yeasts” or “true yeasts.” Other common fungal clades were Basidiomycota:Tremellomycetes (8%), Zygomycota:Mucoromycotina (6%), Ascomycota:Dothideomycetes (6%) and Basidiomycota:Microbotryomycetes (5%).

The 20 most abundant bacterial and fungal OTUs across all butterflies (excluding those detected in less than 10% of frugivores or nectivores) are listed in Tables 2 and 3. The predominant bacteria included known associates of butterflies (*Orbus* spp.; Kim et al 2013; Hammer et al 2014) and other insects with sugar-rich diets (e.g. *Erwinia* sp., *Asaia* sp., *Commensalibacter intestini;* Crotti et al 2010), putative insect parasites and pathogens (e.g. *Spiroplasma* sp., *Serratia* spp., *Wolbachia* sp.), common gut inhabitants (e.g. *Klebsiella*/*Enterobacter, Vagococcus/Enterococcus*), and common fermenting bacteria (e.g. *Lactococcus* sp., *Acetobacter* sp.). The most abundant fungi were largely ascomycotous yeasts (especially members of the genera *Hanseniaspora*, *Pichia*, *Kazachstania* and *Candida*), many of which are known to associate with insects and/or plants— particularly nectars, decaying fruits, and beetle guts (Suh et al 2005; Kurtzman et al 2011). One OTU was most closely identified as a *Rhizopus* species; this genus is commonly associated with decaying vegetable matter (Kirk et al 2008). Likely plant pathogens (e.g. *Clavariopsis* sp. and a Letiomycetes species) and plant-associated basidiomycotous yeasts (OTUs that aligned most closely to the genera *Bensingtonia*) were also common.

Several of the most abundant OTUs (those with the greatest number of counts across all butterflies) were also the most prevalent (detected in the largest number of butterflies). The most frequently detected bacterial OTUs were a *Vagococcus/Enterococcus* species found in 83% of the butterflies, a *Klebsiella/Enterobacter* species found in 82%, and an *Orbus* species found in 73%. The most prevalent fungal OTUs were *Hanseniaspora uvarum* found in 45%, *Hanseniaspora opuntiae* found in 42%, and *Hanseniaspora guilliermondii* found in 40%.

#### 1a. Total bacterial load in the butterfly gut

We quantified the total abundance of bacteria in the guts of 261 butterflies. The total number of 16S rRNA copies in a butterfly’s gut ranged from 5×10^5^ to 1×10^11^, with a median of 7.5×10^8^ (interquartile range: 1.2×10^8^ – 2.7×10^9^). Larger butterfly individuals hosted greater numbers of bacteria (Supplementary Figure 1). Bacterial load differed among host species (Figure 1; random host species intercept term: *df* = 1, χ^2^ = 17.5, *p* < 0.001) but did not differ between feeding guilds after accounting for host species and size (fixed diet term: *df* = 1, χ^2^= 1.8, *p* = 0.18).

**Figure 1.**
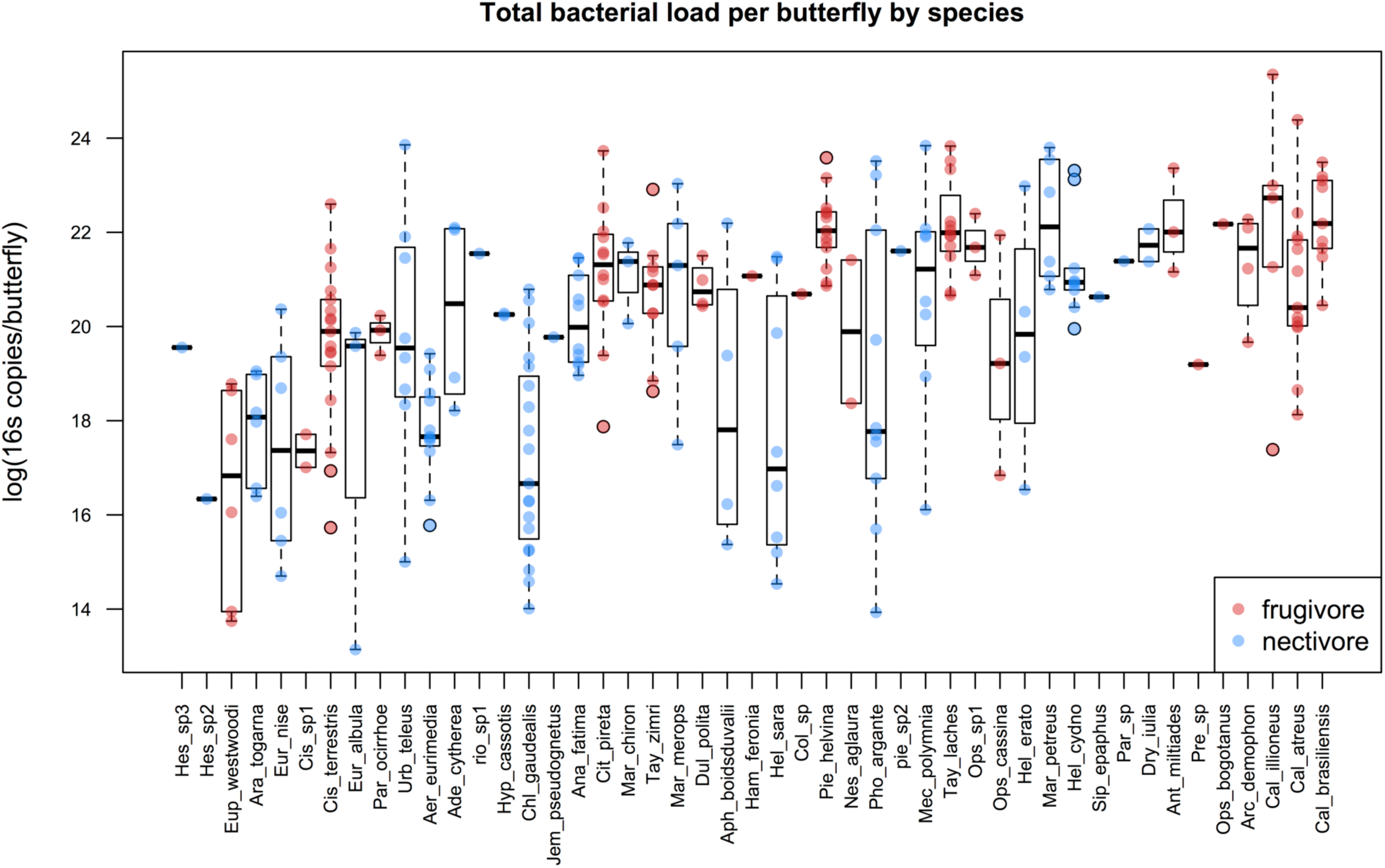
Total bacterial load per butterfly by species. Total bacterial 16s counts per butterfly were estimated via qPCR. Host species are arranged in ascending size (wing length) along the x-axis. Dots indicate copy number for individual frugivores (red) or nectivores (blue), and boxplots depict medians and interquartile ranges of the data. Whiskers are placed at 1.5 times the interquartile range or, if all data fall within this range, they are placed at most extreme value measured. Note that although feeding guild is displayed, it was not a significant predictor of total bacterial load.

### 2. Variation in gut microbial community composition between host dietary guilds, species, and individuals

#### 2a. Variation between the feeding guilds

The gut flora of frugivores and nectivores did not differ from each other in observed OTU richness (Figure 2; bacteria fixed feeding guild term: *df* = 1, χ^2^ = 0.98, p = 0.32; fungi fixed feeding guild term: *df* = 1, χ^2^ = 0.09, p = 0.76). However, gut microbial community composition did differ between the feeding guilds (bacteria: permanova, F= 13.2, R^2^= 0.040, p=0.001, Figure 3a; fungi: permanova, F= 7.4, R^2^= 0.038, p=0.001, Figure 3b). Host feeding guild explained 4% of the variation in both bacterial and fungal community compositions.

**Figure 2.**
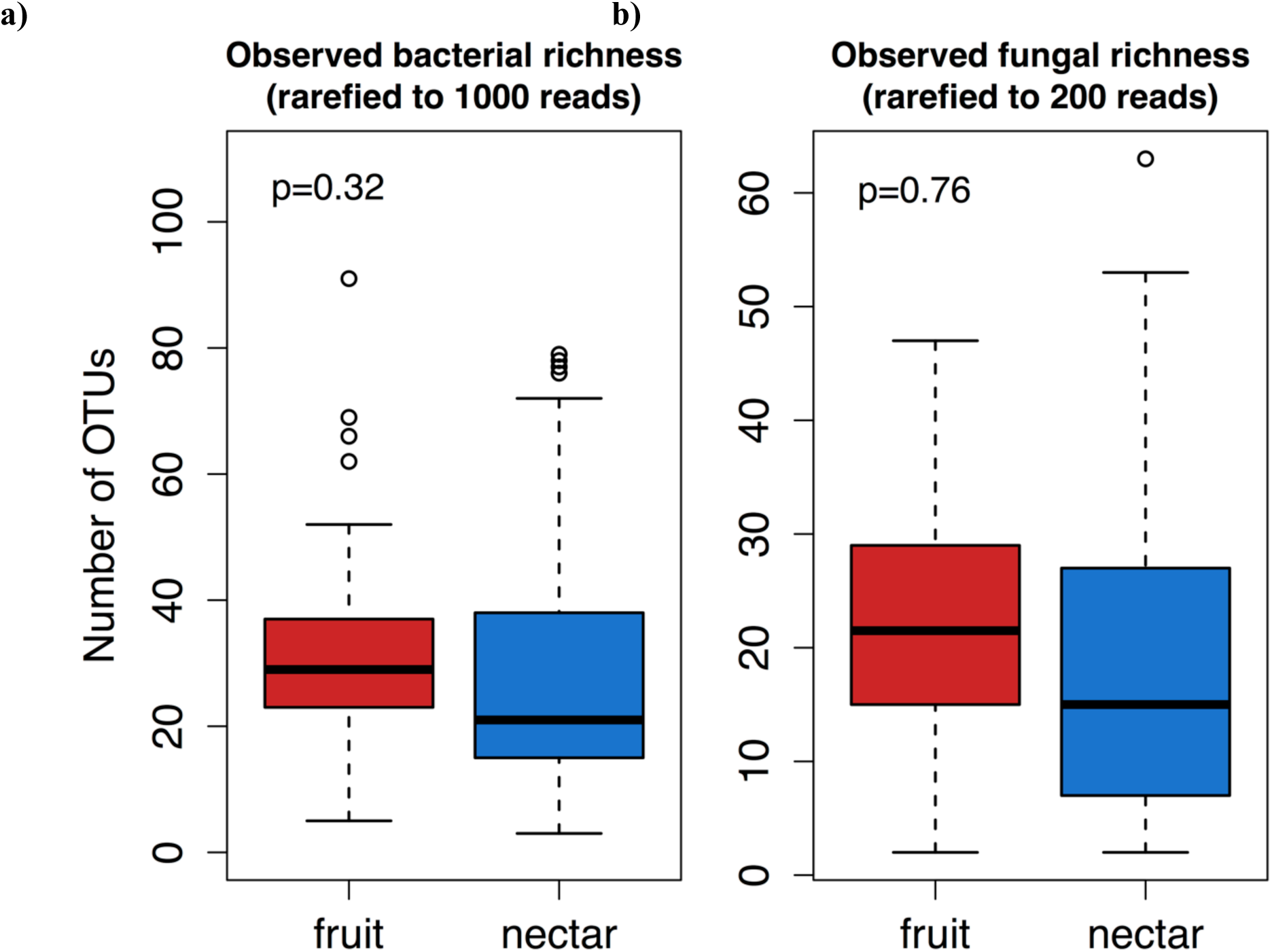
Microbial species richness in frugivores and nectivores. Bacterial (a) and fungal (b) observed species richness per butterfly did not differ between frugivores and nectivores. P-values are the result of model comparisons and control for host species (see text). Boxplot features are as described in Figure 1. One outlier of 162 bacterial OTUs in a nectivore is not shown in panel (a). Note that since bacteria and fungi were rarefied to different cutoffs, species richness estimates are not comparable between the two.

**Figure 3.**
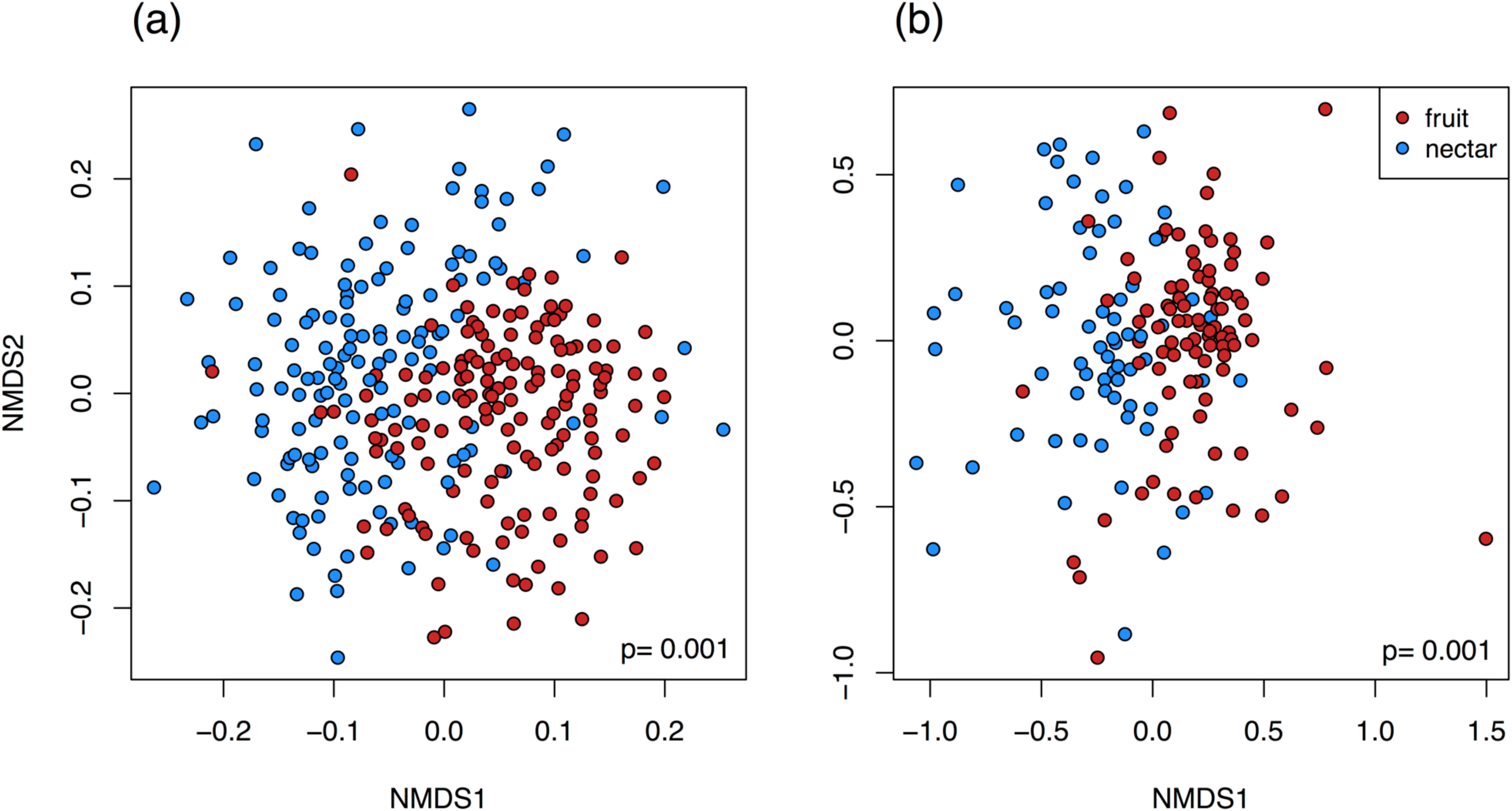
Ordinations of bacterial and fungal community composition. NMDS plots of the Bray-Curtis dissimilarities between butterflies. PerMANOVA tests confirmed that the gut flora of frugivores and nectivores differed in bacterial and fungal OTU composition (p-values displayed on plots). (a) Bacterial gut flora of 148 frugivores and 142 nectivores. (b) Fungal gut flora of 92 frugivores and 69 nectivores. One outlier is not displayed in panel (b).

Despite substantial inter-individual and inter-specific variation in microbial community composition (described below), average relative abundances of 10 of the 20 most abundant bacteria (Figure 4) and fungi (Figure 5) systematically differed between the feeding guilds. Of the bacteria, *Swaminanthia/Asaia* sp., *Bartonella* sp. and a strain of *Commensalibacter intestini* were at higher relative abundance in nectivores than frugivores. Another strain of *Commensalibacter intestini*, *Wolbachia* sp., a Porphyromonadaceae species, *Gilliamella/Orbus* sp., *Orbus* sp. and *Acetobacter* sp. were at higher relative abundance in frugivores than nectivores. Of the fungi, an unidentified fungus (OTU 7), a Letiomycetes species and a Pleosporales species were more abundant in nectivores, while an unidentified ascomycete (OTU 20), two strains of *Kazachstania exigua*, *Hanseniaspora guilliermondii*, *H. uvarum*, *H. opuntiae*, and *Pichia fermentans* associated more strongly with frugivores.

**Figure 4.**
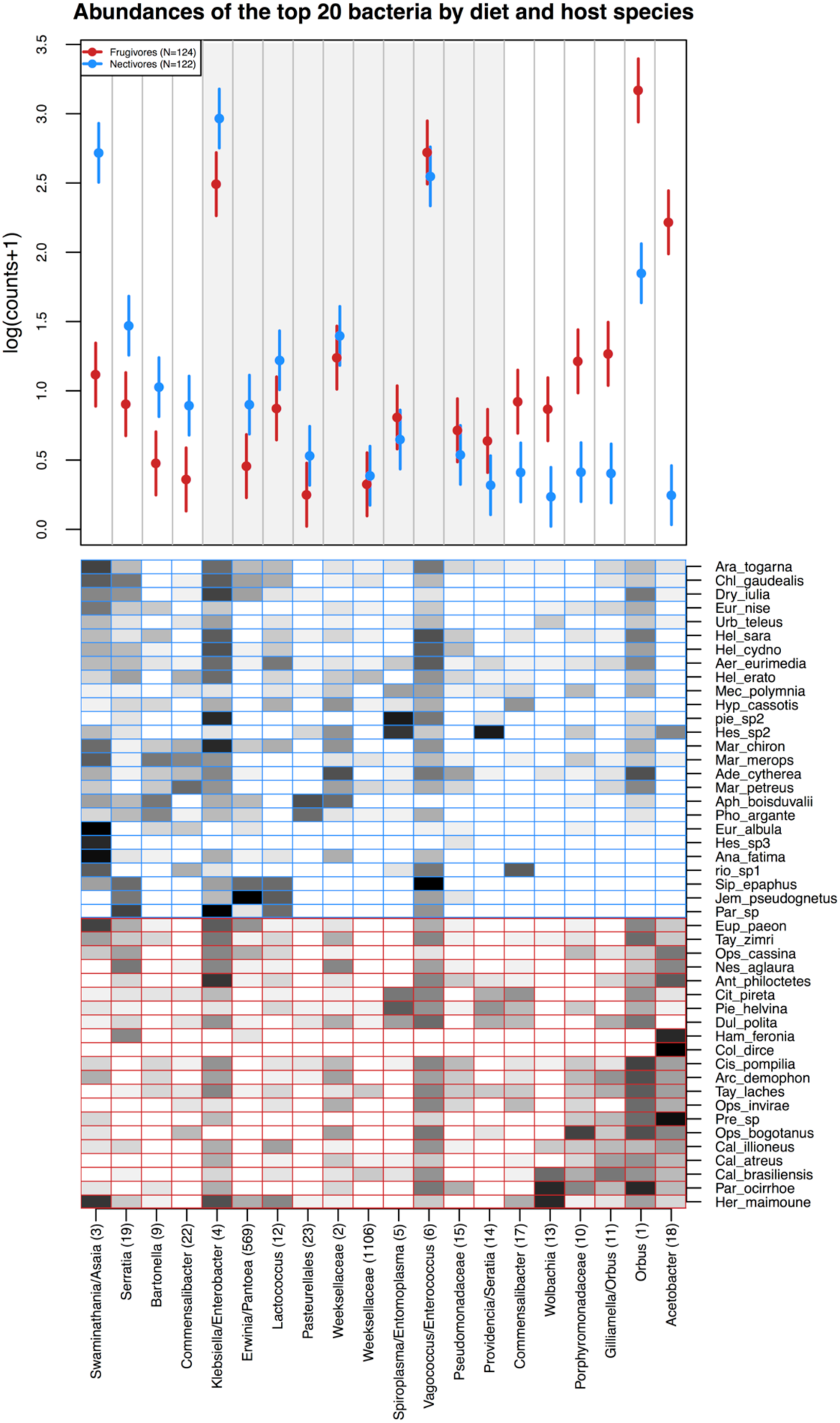
Effect of host diet (a) and host species (b) on bacterial community composition. **(a)** Model-predicted estimates of the mean relative abundance per butterfly (counts out of 1000) of the 20 most abundant bacterial OTUs in the dataset. Relative abundance was modeled as a function of the interaction between host diet and OTU identity, with a random intercept of host species. Points indicate the model estimated mean and lines indicate one standard error. OTUs are arranged along the x-axis according to the magnitude and direction of the difference in their abundance between frugivores and nectivores. Each OTU is designated by genus or by the finest taxonomic resolution available, followed by OTU number in parentheses. The shaded gray box indicates OTUs that did not differ in abundance between the feeding guilds. **(b)** Mean relative abundances of the 20 most abundant bacterial OTUs per host species. Darker shading indicates higher mean relative abundance.

**Figure 5.**
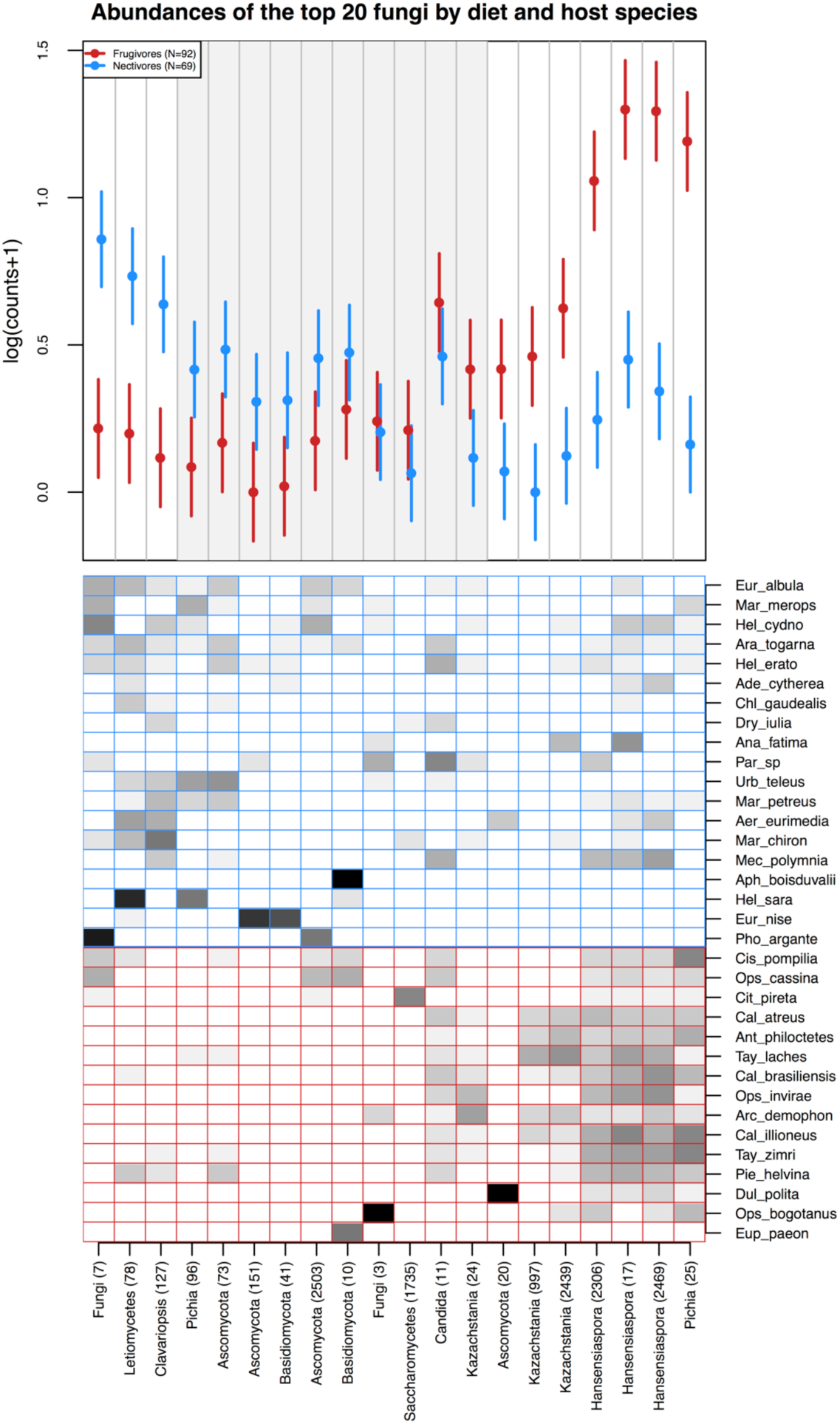
Effect of host diet (a) and host species (b) on fungal community composition. **(a)** Model-predicted estimates of the mean relative abundance per butterfly (counts out of 1000) of the 20 most abundant fungal OTUs in the dataset. Relative abundance was modeled as a function of the interaction between host diet and OTU identity, with a random effect of host species. Points indicate the predicted mean and lines indicate one standard error. OTUs are arranged along the x-axis according to the magnitude and direction of the difference in their abundance between frugivores and nectivores. Each OTU is designated by genus or by the finest taxonomic resolution available, followed by OTU number in parentheses. The shaded gray box indicates OTUs that did not differ in abundance between the feeding guilds. **(b)** Mean relative abundances of the 20 most abundant fungal OTUs per host species. Darker shading indicates higher mean relative abundance.

#### 2b. Variation among host species

Observed OTU richness differed among butterfly species (Supplementary Figure 2; bacteria random host species intercept term: *df* = 1, χ^2^ = 43.3, p < 0.001; fungi random host species intercept term: *df* = 1, χ^2^ = 52.3, p < 0.001). Gut microbial community composition also differed among host species (bacteria: permanova host species term, *df* =35, F= 2.3, R^2^= 0.244, p< 0.001; fungi: permanova host species term, F= 2.3, R^2^= 0.319, p< 0.001). Host species explained 24% and 32% of the variation in bacterial and fungal community composition, respectively. Gut flora were more similar within than between species (Figure 6).

**Figure 6.**
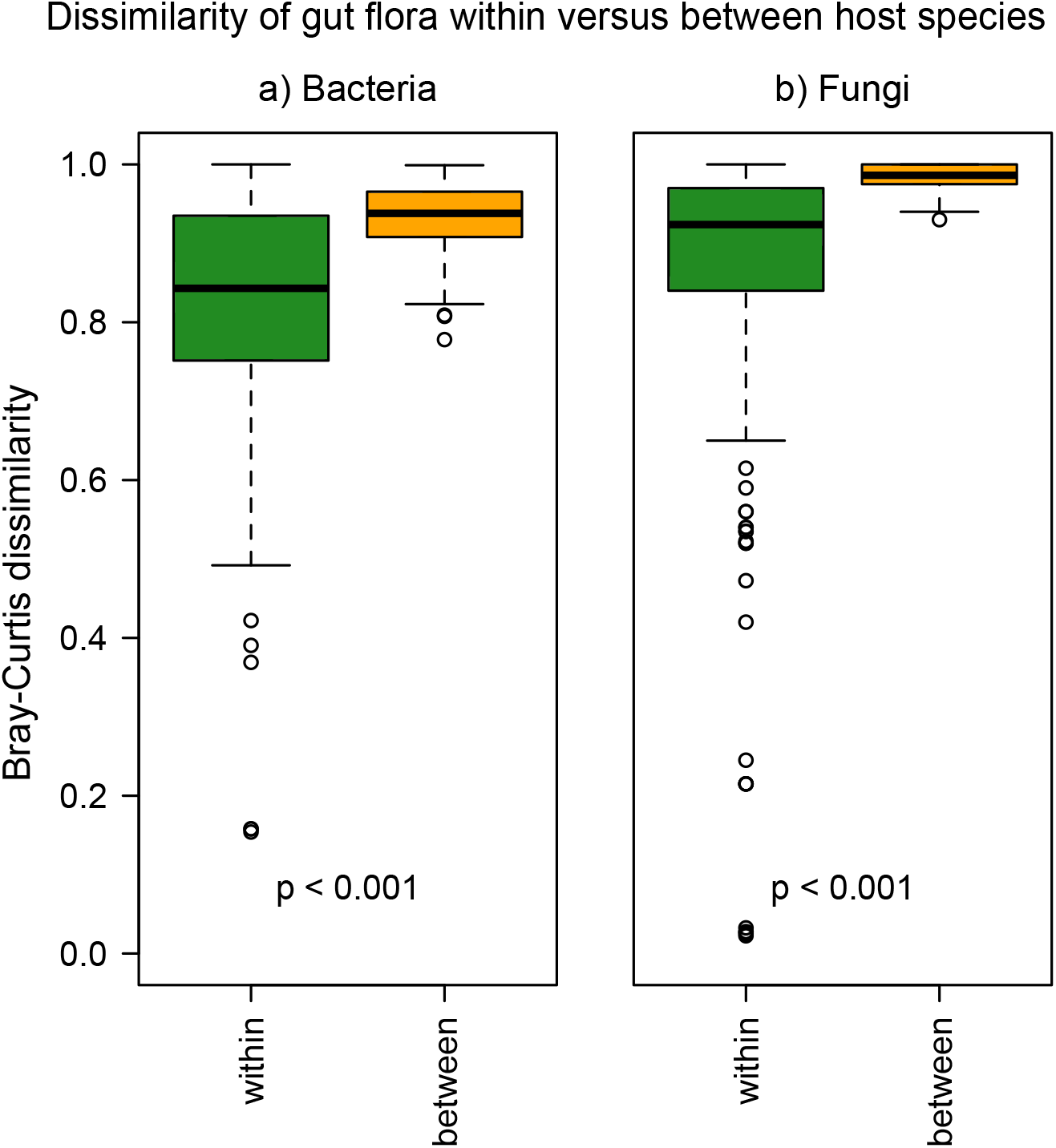
Dissimilarity of gut flora within versus between host species. Every point represents the average Bray-Curtis dissimilarity between of a given individual’s gut community and the gut communities of all other conspecific hosts (green) or all heterospecific hosts (orange). Boxplot features are as described in Figure 1. P-values are the results of t-tests for pairwise differences in dissimilarity.

#### 2c. Variation among individuals

The vast majority of variation in community composition was expressed among individuals: After accounting for species- and guild-level differences, residual variation in community composition was 71.7% for bacteria and 64.4% for fungi (permanova). On average, a pair of butterflies shared 23% (IQR 13%-31%) of their bacterial OTUs and 10% (IQR 0%-16%) of their fungal OTUs. No OTU was present in all butterfly individuals. Each bacterial OTU was found in a mean of 9 individuals (IQR: 1-6) and each fungal OTU was found in a mean of 4 individuals (IQR: 1-3).

### 3. Comparison of the gut flora to food microbial communities

We sequenced the bacterial flora of 87 food samples, 83 of which were retained after rarefaction, and the fungal flora of 75 foods, 66 of which were retained after rarefaction (Table 1; Supplementary Table 2). After sequence processing and quality filtering we obtained 7.7 million bacterial and 3.2 fungal sequences from potential food sources (fruits, nectars, and trap baits). After rarefying, a total of 1205 bacterial OTUs and 529 fungal OTUs were observed across all the food samples. (Again note that numbers of bacterial and fungal OTUs cannot be compared due to differences in rarefaction depth.)

The microbial communities in wild fruit juice, the fermented trap baits, and wild nectars were distinct from each other and from butterfly gut communities (Figure 7; Figure 8). In fact, butterfly gut communities were highly dissimilar to the microbial communities on their food sources. For example, frugivores and wild fruits shared 7% of their bacterial OTUs on average (IQR 3%-9%), and the median Bray-Curtis dissimilarity between their bacterial communities was 0.91 (Figure 8a). Despite these pronounced differences, frugivores’ gut bacteria more closely resembled the microbial composition of the trap baits than nectivores’ gut bacteria did, and similarly, nectivores’ gut communities were more similar to nectar bacteria than frugivores’ gut flora were (Figure 8a). Frugivores’ and nectivores’ bacterial communities, however, were equivalently distinct from those inhabiting the surfaces of wild fruits (Figure 8a).

**Figure 7.**
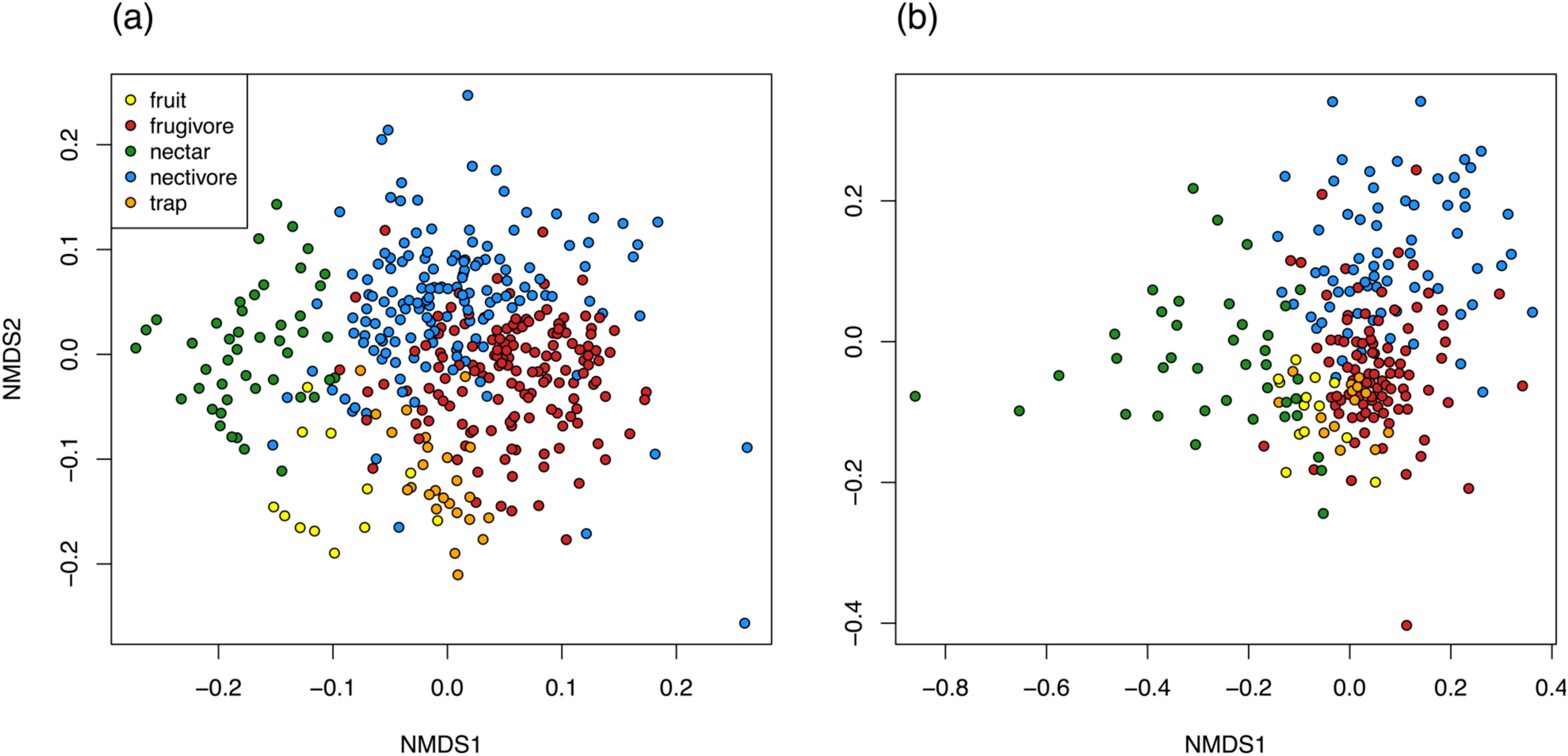
Comparison of frugivore and nectivore gut communities to the microbial communities of butterfly foods. NMDS plots of the Bray-Curtis dissimilarities between butterflies and their foods. (a) Bacterial flora of 149 frugivores, 149 nectivores, 46 nectars, 25 trap baits, and 12 fruits. (b) Fungal gut flora of 94 frugivores, 70 nectivores, 39 nectars, 14 trap baits, and 13 fruits. Two samples were fungal outliers and are not displayed in panel (b). (Sample sizes differ slightly from Figure 3 because in order to include as many food samples as possible, the combined gut and food sequence data were rarefied to different depths than the gut sequence data alone.)

**Figure 8.**
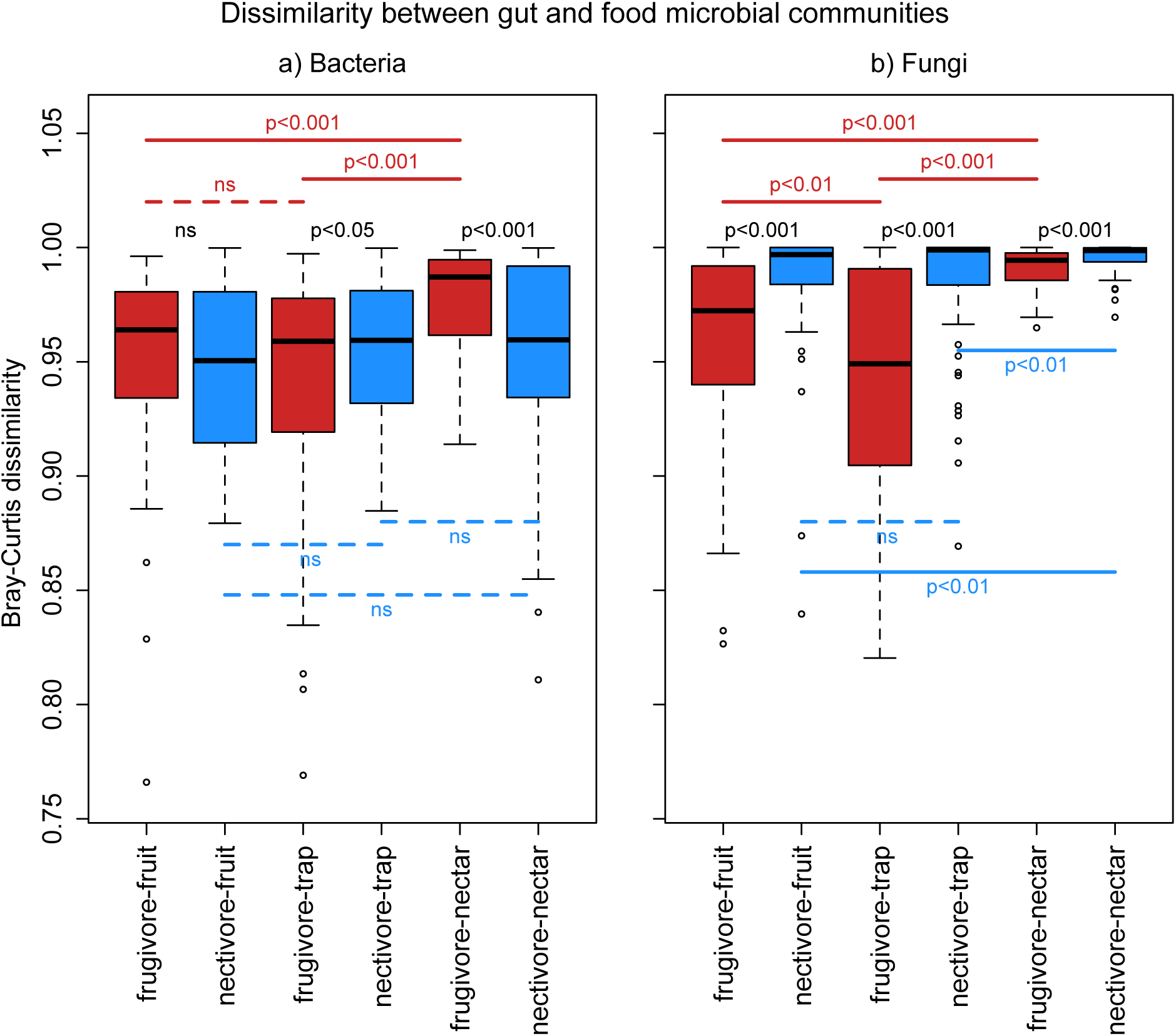
Dissimilarity between gut and food microbial communities. Bray-Curtis dissimilarities between butterfly gut bacterial (a) and fungal (b) communities and the microbial communities in butterfly foods. Every point represents the average distance of a butterfly gut community to all food communities in a given category (fruit, trap bait, or nectar). Comparisons to frugivorous butterflies are colored red; comparisons to nectivores are colored blue. Boxplot features are as described in Figure 1. P-values are the results of t-tests for pairwise differences in dissimilarity; these have been FDR-corrected. Note that three fungal strains of *Kazachstania exigua* were excluded from the butterfly data but not the food data, since these OTUs were introduced into frugivores’ guts via the trap baits. (Described in the main text.) Frugivores would appear more similar to the trap baits if these OTUs had been included.

Fungal communities in frugivore guts more closely resembled those of wild fruits and trap baits than did those of nectivore guts, as expected (Figure 8b). Surprisingly, frugivores’ gut fungal communities were more similar, on average, to fungi in nectar than nectivores’ gut communities were (Figure 8b), though the effect size was extremely small.

While gut communities were distinct from food microbial flora in terms of species composition and relative abundances, the percentage of bacterial gut reads assigned to OTUs that were also present in the host’s diet was generally high. On average, about 85% percent of gut bacterial reads derived from OTUs present in their host’s food. Fungal OTUs present in food, however, accounted for a comparatively small 24% to 40% of butterfly gut fungal reads.

### 4. Functional abilities: gut microbial community catabolism

We measured the catabolic profiles of 248 gut communities via culture-based assays. Gut communities digested an average of 51 substrates (IQR: 41-63) out of 71 total. As two communities diverged in bacterial species composition, their catabolic profiles also became more different; however, the effect size was very small (Mantel test: r = 0.067, p = 0.013). Catabolism was not correlated with fungal community composition (Mantel test: p = 0.50).

Across the entire dataset, host diet explained just 3% of the total variation in community catabolic profile (permanova on Euclidean distances between catabolic profiles; *df* =1, F= 7.6, R^2^=0.029, p=0.001), while host species identity explained 18% of the variation (*df*= 33, F= 1.4, R^2^=0.180, p=0.001). After accounting for diet- and species-level differences, residual variation among individuals in catabolic profile was 79.1%.

Lack of strong correlation between catabolic profile and host diet could be due to the fact that not all catabolic functions are relevant for bacteria in butterfly guts. To investigate diet-based differences in catabolic profile in more detail, we modeled catabolism as a function of host diet, substrate identity, and substrate class, controlling for butterfly identity and number of 16S rRNA copies in the butterfly. We also tested an alternate random effects structure, nesting butterfly individuals within butterfly species, but this did not improve model fit (anova: *df* = 1, χ^2^= 0.258 p =0.61). Catabolism of several individual substrates differed markedly between the feeding guilds (feeding guild by substrate interaction term: *df* = 70, χ^2^ = 428.9 p <<0.001). Frugivorous gut flora catabolized 15 substrates more actively than nectivorous gut communities did, with the most marked difference being in catabolism of D-serine, saccharic acid, mucic acid, L-serine and lactic acid (Supplementary Table 5). Nectivorous gut flora digested 10 substrates more actively, with the most extreme differences in catabolism of mannitol, sucrose, glucose, fructose, and maltose (Supplementary Table 5). Entire classes of nutrients were also catabolized differently between the guilds (feeding guild by substrate class interaction term: *df* = 6, χ^2^ = 61.7, p <<0.001). Specifically, nectivores’ gut flora catabolized sugars and sugar alcohols more actively than frugivorous gut flora, while frugivores’ gut flora were more successful at digesting amino acids, carboxylic acids, and dicarboxylic acids (Figure 9; Supplementary Table 6).

**Figure 9.**
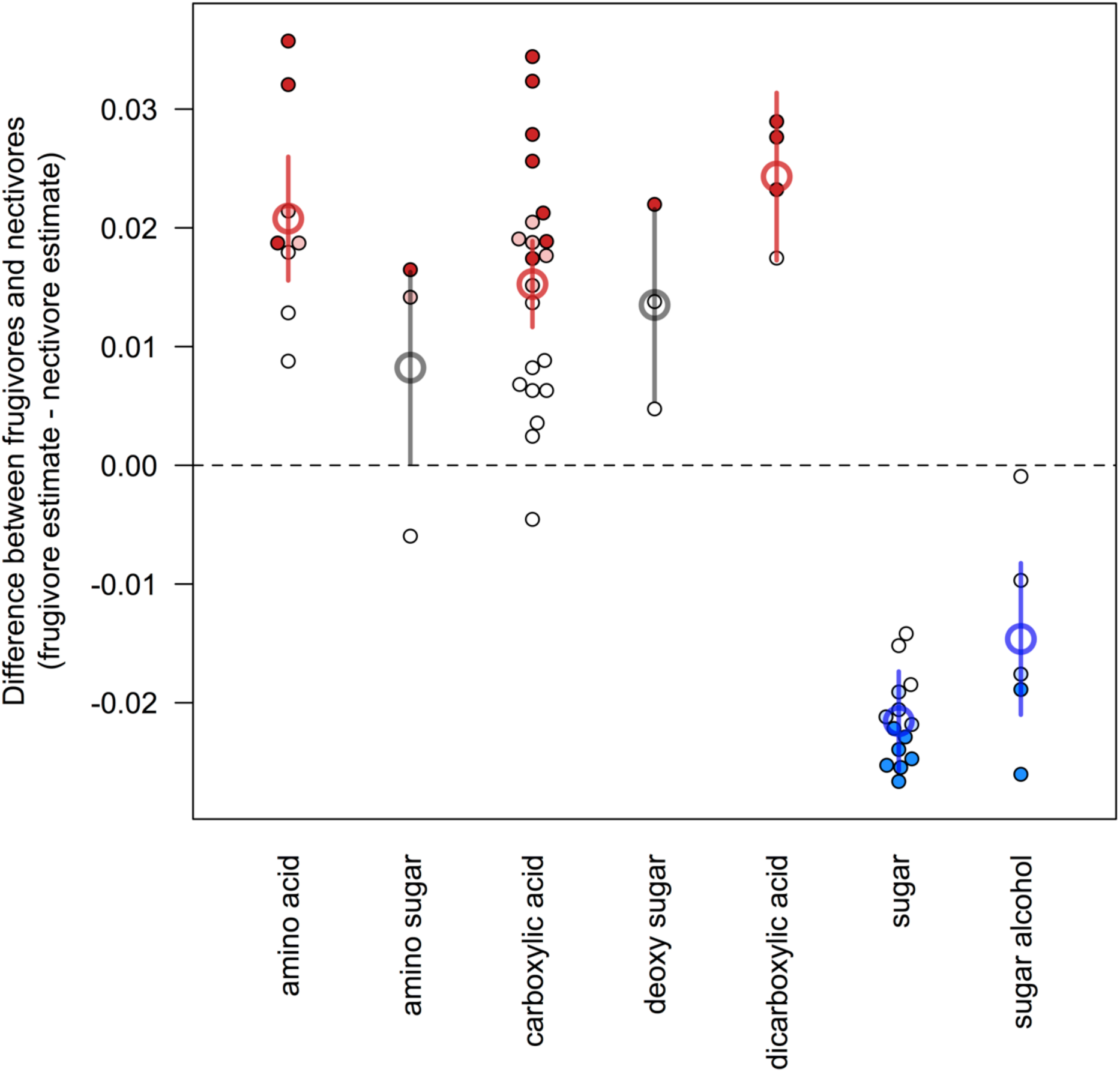
Differences in microbial community catabolism between frugivores and nectivores. Points indicate the model-estimated mean difference in catabolism of a substrate (units of absorbance at 590 nm, standardized by plate and Hellinger transformed) between frugivores’ and nectivores’ gut microbial communities. Gut community catabolism of substrates in dark red and dark blue was significantly greater in frugivores or nectivores, respectively, after multiple test correction. Light red and light blue substrates were significantly different between guilds before, but not after multiple test correction. Thick lines and points indicate model-estimated means and standard errors for substrate classes. Red indicates classes that were catabolized more strongly by frugivore gut communities; blue indicates classes that were digested more actively by nectivore gut flora.

Total number of 16S rRNA copies was not significantly correlated with catabolism (p= 0.09 for the model of substrate identity and p=0.07 for the model of substrate class), suggesting that our standardization procedure adequately controlled for any initial differences in cell inoculation densities.

## Discussion

We evaluated how microbial community structure and catabolic function varied among host feeding guilds, species and individuals within a community of over 50 species of Neotropical butterflies. On average, 30 bacterial OTUs and 21 fungal OTUs were detected per gut. Although the bacterial and fungal communities in adult butterfly guts varied substantially among individual hosts, they did differentiate among host species and between host feeding guilds. Gut community composition varied less between host feeding guilds than among host species, suggesting that other non-dietary aspects of host biology play a large role in structuring the butterfly gut flora. Gut communities were distinct in composition from the microbial flora found in butterfly foods, indicating that the adult butterfly gut environment strongly filters potential colonists. Despite high variability in the gut microbial community, host dietary guild was nevertheless associated with consistent changes in both the relative abundances of dominant microbes and the catabolic capacities of the gut flora. Frugivorous gut communities were better at digesting amino acids and carboxylic acids, while nectivorous gut flora outperformed frugivorous communities in catabolism of sugars and sugar alcohols. These catabolic patterns are congruent with the relative nutritional makeup of butterfly foods (Ravenscraft and Boggs 2016), which suggests that gut flora specialize in digestion of compounds abundant in the host’s diet.

### 1. Microbes present in the guts of adult butterflies

#### 1a. Bacteria

The composition of the butterfly’s bacterial flora is broadly similar to that of other insects: at the phylum-level, Proteobacteria comprised 68% and Firmicutes 12% of all reads while in comparison, a meta-analysis of the insect gut flora reported that 57% were Proteobacteria and 22% were Firmicutes (Colman et al 2012). At finer taxonomic resolution (see Table 2), the most abundant bacterial OTUs were mostly known gut colonists, particularly those associated with sugar-rich diets, and OTUs related to insect parasites and pathogens. (Individual OTUs are discussed in more detail below.) Total bacterial load varied among species, but there were no consistent differences between the feeding guilds after controlling for differences in size among individual hosts. Larger butterfly individuals hosted greater numbers of bacteria. This conforms previous work which demonstrated that total microbial load scales with host size (Kieft and Simmons 2015).

**Table 2.** Taxonomic identities of the 20 most abundant bacterial OTUs. The 20 most abundant bacterial OTUs. “% Total” is the percentage of sequences (pooled across all butterflies) that were assigned to the OTU. “% Butterflies”, “% Frugivores”, and “% Nectivores” are the percentages of all butterflies, frugivores, and nectivores in which the OTU was detected. Taxonomy was assigned by the RDP classifier using the Greengenes training set, except as otherwise noted. Confidences measure the degree of certainty of the taxonomic assignment.

| OTU ID | % Reads | % Butterflies | % Frugivores | % Nectivores | Frug:Nect | Phylum | Class | Order | Family | Genus | Species | Confidence | Notes |
| --- | --- | --- | --- | --- | --- | --- | --- | --- | --- | --- | --- | --- | --- |
| OTU_1 | 9.8 | 73 | 84 | 63 | 1.3 | Proteobacteria | Gammaproteobacteria | Orbales | Orbaceae | *Orbus* |  | 100 | a |
| OTU_4 | 8.7 | 82 | 81 | 84 | 1.0 | Proteobacteria | Gammaproteobacteria | Enterobacteriales | Enterobacteriaceae | *Klebsiella/Enterobacter* |  | 100 | b |
| OTU_3 | 8.3 | 56 | 42 | 70 | 0.6 | Proteobacteria | Alphaproteobacteria | Rhodospirillales | Acetobacteraceae | *Swaminathania/Asaia* |  | 97 | c |
| OTU_6 | 6.0 | 83 | 86 | 80 | 1.1 | Firmicutes | Bacilli | Lactobacillales | Enterococcaceae | *Vagococcus/Enterococcus* |  | 62 | d |
| OTU_2 | 4.8 | 44 | 41 | 48 | 0.9 | Bacteroidetes | Flavobacteriia | Flavobacteriales | [Weeksellaceae] |  |  | 96 |  |
| OTU_19 | 3.2 | 46 | 34 | 59 | 0.6 | Proteobacteria | Gammaproteobacteria | Enterobacteriales | Enterobacteriaceae | *Serratia* | *marcescens* | 94 |  |
| OTU_5 | 2.7 | 28 | 27 | 29 | 0.9 | Tenericutes | Mollicutes | Entomoplasmatales |  | *Spiroplasma/Entomoplasma* |  | 100 | e |
| OTU_10 | 2.6 | 28 | 36 | 20 | 1.8 | Bacteroidetes | Bacteroidia | Bacteroidales | Porphyromonadaceae |  |  | 68 |  |
| OTU_9 | 2.5 | 30 | 27 | 34 | 0.8 | Proteobacteria | Alphaproteobacteria | Rhizobiales | Bartonellaceae | *Bartonella* |  | 92 | f |
| OTU_12 | 2.5 | 44 | 40 | 49 | 0.8 | Firmicutes | Bacilli | Lactobacillales | Streptococcaceae | *Lactococcus* |  | 100 |  |
| OTU_18 | 2.3 | 46 | 72 | 19 | 3.8 | Proteobacteria | Alphaproteobacteria | Rhodospirillales | Acetobacteraceae | *Acetobacter* |  | 100 | g |
| OTU_11 | 2.2 | 34 | 46 | 22 | 2.1 | Proteobacteria | Gammaproteobacteria | Orbales | Orbaceae | *Gilliamella/Orbus* |  | 91 | h |
| OTU_17 | 2.2 | 27 | 36 | 18 | 2.0 | Proteobacteria | Alphaproteobacteria | Rhodospirillales | Acetobacteraceae | *Commensalibacter* | *intestini* | 95 |  |
| OTU_14 | 1.7 | 19 | 25 | 13 | 1.9 | Proteobacteria | Gammaproteobacteria | Enterobacteriales | Enterobacteriaceae | *Providencia/Serratia* |  | 73 | i |
| OTU_13 | 1.7 | 18 | 22 | 13 | 1.7 | Proteobacteria | Alphaproteobacteria | Rickettsiales | Rickettsiaceae | *Wolbachia* |  | 100 |  |
| OTU_15 | 1.6 | 26 | 31 | 20 | 1.5 | Proteobacteria | Gammaproteobacteria | Pseudomonadales | Pseudomonadaceae |  |  | 73 |  |
| OTU_23 | 1.4 | 23 | 26 | 19 | 1.4 | Proteobacteria | Gammaproteobacteria | Pasteurellales |  |  |  | 100 |  |
| OTU_569 | 1.4 | 27 | 18 | 37 | 0.5 | Proteobacteria | Gammaproteobacteria | Enterobacteriales | Enterobacteriaceae | *Erwinia/Pantoea* |  | 56 | j |
| OTU_22 | 1.4 | 25 | 23 | 27 | 0.9 | Proteobacteria | Alphaproteobacteria | Rhodospirillales | Acetobacteraceae | *Commensalibacter* | *intestini* | 93 |  |
| OTU_1106 | 1.2 | 19 | 16 | 21 | 0.8 | Bacteroidetes | Flavobacteriia | Flavobacteriales | [Weeksellaceae] |  |  | 68 |  |

**Table 3.**
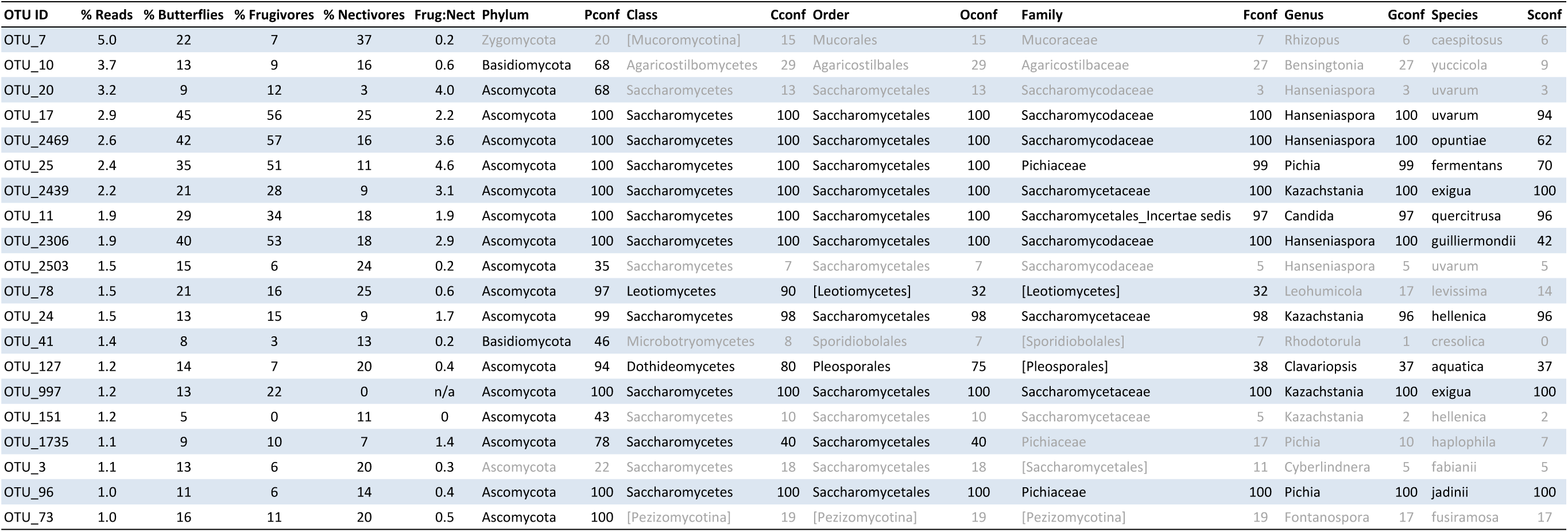
Taxonomic identities of the 20 most abundant fungal OTUs. The 20 abundant fungal OTUs detected in at least 10% of the butterfly samples. “% Total” is the percentage of sequences (pooled across all butterflies) that were assigned to the OTU. “% Butterflies”, “% Frugivores”, and “% Nectivores” are the percentages of all butterflies, frugivores, and nectivores in which the OTU was detected. Taxonomy was assigned by the RDP classifier using the Warcup training set. Where a rank is *incertae sedis*, the closest available rank is written in brackets. Confidences measure the degree of certainty of the taxonomic assignment and are cumulative from higher to lower taxonomic ranks. Assignments with particularly low confidence are greyed out. (One additional fungal OTU, a *Torulaspora* species, was observed at high abundance in the pooled data but was detected in less than 10% of frugivores and less than 10% of nectivores, so is omitted here.)

| OTU ID | % Reads | % Butterflies | % Frugivores | % Nectivores | Frug:Nect | Phylum | Pconf | Class | Cconf | Order | Oconf | Family | Fconf | Genus | Gconf | Species | Sconf |
| --- | --- | --- | --- | --- | --- | --- | --- | --- | --- | --- | --- | --- | --- | --- | --- | --- | --- |
| OTU_7 | 5.0 | 22 | 7 | 37 | 0.2 | Zygomycota | 20 | [Mucoromycotina] | 15 | Mucorales | 15 | Mucoraceae | 7 | Rhizopus | 6 | caespitosus | 6 |
| OTU_10 | 3.7 | 13 | 9 | 16 | 0.6 | Basidiomycota | 68 | Agaricostilbomycetes | 29 | Agaricostilbales | 29 | Agaricostilbaceae | 27 | Bensingtonia | 27 | yuccicola | 9 |
| OTU_20 | 3.2 | 9 | 12 | 3 | 4.0 | Ascomycota | 68 | Saccharomycetes | 13 | Saccharomycetales | 13 | Saccharomycodaceae | 3 | Hanseniaspora | 3 | uvarum | 3 |
| OTU_17 | 2.9 | 45 | 56 | 25 | 2.2 | Ascomycota | 100 | Saccharomycetes | 100 | Saccharomycetales | 100 | Saccharomycodaceae | 100 | Hanseniaspora | 100 | uvarum | 94 |
| OTU_2469 | 2.6 | 42 | 57 | 16 | 3.6 | Ascomycota | 100 | Saccharomycetes | 100 | Saccharomycetales | 100 | Saccharomycodaceae | 100 | Hanseniaspora | 100 | opuntiae | 62 |
| OTU_25 | 2.4 | 35 | 51 | 11 | 4.6 | Ascomycota | 100 | Saccharomycetes | 100 | Saccharomycetales | 100 | Pichiaceae | 99 | Pichia | 99 | fermentans | 70 |
| OTU_2439 | 2.2 | 21 | 28 | 9 | 3.1 | Ascomycota | 100 | Saccharomycetes | 100 | Saccharomycetales | 100 | Saccharomycetaceae | 100 | Kazachstania | 100 | exigua | 100 |
| OTU_11 | 1.9 | 29 | 34 | 18 | 1.9 | Ascomycota | 100 | Saccharomycetes | 100 | Saccharomycetales | 100 | Saccharomycetales_Incertae sedis | 97 | Candida | 97 | quercitrusa | 96 |
| OTU_2306 | 1.9 | 40 | 53 | 18 | 2.9 | Ascomycota | 100 | Saccharomycetes | 100 | Saccharomycetales | 100 | Saccharomycodaceae | 100 | Hanseniaspora | 100 | guilliermondii | 42 |
| OTU_2503 | 1.5 | 15 | 6 | 24 | 0.2 | Ascomycota | 35 | Saccharomycetes | 7 | Saccharomycetales | 7 | Saccharomycodaceae | 5 | Hanseniaspora | 5 | uvarum | 5 |
| OTU_78 | 1.5 | 21 | 16 | 25 | 0.6 | Ascomycota | 97 | Leotiomycetes | 90 | [Leotiomycetes] | 32 | [Leotiomycetes] | 32 | Leohumicola | 17 | levissima | 14 |
| OTU_24 | 1.5 | 13 | 15 | 9 | 1.7 | Ascomycota | 99 | Saccharomycetes | 98 | Saccharomycetales | 98 | Saccharomycetaceae | 98 | Kazachstania | 96 | hellenica | 96 |
| OTU_41 | 1.4 | 8 | 3 | 13 | 0.2 | Basidiomycota | 46 | Microbotryomycetes | 8 | Sporidiobolales | 7 | [Sporidiobolales] | 7 | Rhodotorula | 1 | cresolica | 0 |
| OTU_127 | 1.2 | 14 | 7 | 20 | 0.4 | Ascomycota | 94 | Dothideomycetes | 80 | Pleosporales | 75 | [Pleosporales] | 38 | Clavariopsis | 37 | aquatica | 37 |
| OTU_997 | 1.2 | 13 | 22 | 0 | n/a | Ascomycota | 100 | Saccharomycetes | 100 | Saccharomycetales | 100 | Saccharomycetaceae | 100 | Kazachstania | 100 | exigua | 100 |
| OTU_151 | 1.2 | 5 | 0 | 11 | 0 | Ascomycota | 43 | Saccharomycetes | 10 | Saccharomycetales | 10 | Saccharomycetaceae | 5 | Kazachstania | 2 | hellenica | 2 |
| OTU_1735 | 1.1 | 9 | 10 | 7 | 1.4 | Ascomycota | 78 | Saccharomycetes | 40 | Saccharomycetales | 40 | Pichiaceae | 17 | Pichia | 10 | haplophila | 7 |
| OTU_3 | 1.1 | 13 | 6 | 20 | 0.3 | Ascomycota | 22 | Saccharomycetes | 18 | Saccharomycetales | 18 | [Saccharomycetales] | 11 | Cyberlindnera | 5 | fabianii | 5 |
| OTU_96 | 1.0 | 11 | 6 | 14 | 0.4 | Ascomycota | 100 | Saccharomycetes | 100 | Saccharomycetales | 100 | Pichiaceae | 100 | Pichia | 100 | jadinii | 100 |
| OTU_73 | 1.0 | 16 | 11 | 20 | 0.5 | Ascomycota | 100 | [Pezizomycotina] | 19 | [Pezizomycotina] | 19 | [Pezizomycotina] | 19 | Fontanospora | 17 | fusiramosa | 17 |

The most abundant gut residents in our data were similar to those found in adult *Heliconius erato* butterflies in Panama (Hammer et al 2014). The genera *Orbus*, *Enterobacter*, *Asaia, Enterococcus*, *Lactococcus* and *Commensalibacter* were abundant in both studies, and *Orbus* has also been isolated from the gut of the butterfly *Sasakia charonda* in South Korea (Kim et al 2013). This suggests that adult butterflies do form consistent associations with some gut microbes. What drives these associations is uncertain. Butterflies might maintain specific relationships with some microbes via mechanisms such as vertical transmission. Gut flora can be transmitted from mother to offspring via internal migration of gut microbes into the eggs, as observed in the moth *Galleria mellonella*, or via the smearing of gut microbes on egg surfaces, which has been hypothesized to occur in tobacco hornworm (*Manduca sexta*) (Brinkmann et al 2008; Freitak et al 2014). The relative frequency of such inter-generational transmission is unclear, however, because the majority of the larval gut flora are purged during metamorphosis (Kingsley 1972; Hammer et al 2014; but see Johnston and Rolff 2015). Other members of the gut flora may be common in both butterflies and other insects because they are generally adapted to the insect gut environment (e.g. Chouaia et al 2012) and opportunistically colonize many species. Many of the most abundant genera we detected may fall into this category.

#### 1b. Gut fungi were primarily yeasts and common plant associates

The most abundant fungal OTUs were common plant and insect associates. Two thirds of all fungal sequences belonged to the order Saccharomycetales, the “budding yeasts” or “true yeasts.” These single-celled fungi are often associated with sugar-rich environments and decaying vegetable matter. Most of the remaining OTUs with reliable taxonomic assignments were likely plant associates or pathogens. Studies characterizing the gut flora often focus on bacteria and overlook fungi, but available evidence suggests that some insect guts host a rich diversity of fungi, especially yeasts (Suh et al 2005). Some of these fungi may play an active role in the gut, while others may simply use insects as a means of dispersal (Blackwell and Jones 1997; Starmer and Lachance 2011).

### 2. Variation in gut microbial community composition between host dietary guilds, among species, and among individuals

#### 2a. A smaller degree of variation was attributable to differences between feeding guilds

Frugivores and nectivores did not differ in total bacterial load nor in bacterial or fungal species richness, but did differ in the composition of their microbial flora. The degree of variation explained by dietary guild was relatively small (about 4% for bacteria and fungi), however it was comparable to that observed in many studies of microbial communities and is likely biologically relevant: relative abundances of half of the most common OTUs differed systematically between the feeding guilds. In general, the fungi more abundant in nectivores were closely related to plant pathogens, while the fungi more abundant in frugivores were yeasts associated with fruit decay, suggesting that this variation likely derived from exposure to different microbial source pools. In contrast, there was no pattern in the types of bacteria that differed in abundance between frugivores and nectivores.

#### 2b. A moderate degree of variation was attributable to differences among host species

About 24% and 32% percent of the variation in bacterial and fungal OTU compositions, respectively, was attributed to differences among host species. Total bacterial load and bacterial and fungal OTU richness also differed among butterfly species. Interspecies variation in the butterfly gut flora could result from behavioral differences that expose butterflies to different microbial pools. Such species traits could include habitat preferences. For example, species that favor the forest understory could be exposed to more soil and wood decay fungi than those that prefer the canopy. Additionally, many, but not all species drink from mud, dung, or carrion, in a behavior known as “puddling” (Adler and Pearson 1982; Sculley and Boggs 1996), and variation in these auxiliary feeding substrates could introduce different microbes to the gut. Differences could also arise from variation in gut chemistry: for example, the guts of larger butterflies might be more anoxic, or there could be interspecies differences in gut pH.

At least some of these factors could exhibit evolutionary signal in butterflies, resulting in correlation between the butterfly phylogeny and gut community composition. Alternatively or additionaly, vertical transmission of a fraction of the gut community from parent to offspring could result in parallel divergence of microbiota and hosts. The potential for such evolutionary patterns to contribute to interspecies differences in gut community composition is beyond the scope of the current study, but will be addressed in a future paper (Ravenscraft et al, in prep).

#### 2c. Variation in microbial community composition among individual butterflies is high, but not unusual

There was a great deal of variation among individual butterflies in microbial community composition: the most prevalent bacterial and fungal OTUs were detected in only 83% and of 45% butterflies, respectively, and half of all OTUs were found in only one individual. Although this degree of variation may seem extreme, it is comparable to that observed in other animals: human colon microbiota were 34% similar among individuals in one study (Green et al 2006), and in another, no OTUs were universally shared and 79% were found in only a single individual (Tap et al 2009). A study of mammalian gut flora found that, on average, 56% of OTUs in an individual mammal were unique to that individual (Ley et al 2008a). It has been proposed that the high degree of variation observed in gut communities is the result of functional redundancy at the OTU level: many microbes may require similar gut environmental conditions and/or serve similar functional roles, leading to communities that differ in OTU membership but not in overall function (Ley et al 2006; Peay et al 2016). However, it is also possible that variation in membership *does* lead to significant variation in function (Peay et al 2016) and that the butterfly gut flora do not play a consistent role across individuals.

#### 2d. The effect sizes of variables related to gut community composition are rarely reported or compared

We had expected that a butterfly’s feeding guild would explain a larger amount of variation in gut community membership than host species identity, but we found the opposite. The relative importance of dietary guild as a determinant of gut community composition has been obscured in the literature by a trend of reporting only the significance (p-value) and not the strength (either the effect size or R-squared) of these relationships. Diet has been shown to significantly affect the community composition of the gut flora in many systems, including mammals (Ley et al 2008a), fish (Sullam et al 2012), and insects (Colman et al 2012), but the strength of this correlation was only reported for the last case. In a broad survey of the insect gut flora, feeding guild (e.g. herbivore, omnivore, detritivore, pollenivore) explained 23% of variation in the gut flora. The comparatively small amount of variation which diet explains in the butterfly gut flora could be due in part to the broadly similar nature of butterfly foods (both being sugary liquids).

### 3. Comparison of gut flora to food microbial communities

#### 3a. Butterfly gut communities were highly dissimilar to fruit and nectar communities, but generally shared more OTUs with the diet of their host than the diet of the other feeding guild

Host diet may shape the gut flora by serving as a source pool of microbes. Since it is likely that adult butterflies acquire a substantial fraction of their gut flora from the environment after emergence from the pupa (Hammer et al 2014), the microbial composition of adult butterfly foods could be especially relevant. We asked whether the gut flora of Neotropical butterflies might derive from their foods, and tested whether differences between frugivore and nectivore gut communities were driven, in part, by differences in the microbes present in fruits and nectars. Butterfly gut communities were more dissimilar to food microbial communities than we had anticipated, with median percentage OTUs shared between a gut community and the microbes residing in its host’s food ranging from 5 to 16% for bacteria and 0 to 14% for fungi. However, gut flora were generally more similar to the community on their host’s food than to microbes on the food of the other feeding guild. This suggests that a limited number of food-derived bacteria and fungi either pass through or colonize the gut. Indeed, comparison of frugivores caught with aerial nets to those caught via baited traps indicated that the yeast *Kazachstania exigua* was transferred from the fermented baits into trapped butterflies’ guts.

The high degree of differentiation between gut and food communities was surprising because the guts of many nonsocial insects do seem to be colonized by environmental bacteria (Ley et al 2008b; Boissière et al 2012). Indeed, prior studies of lepidopterans have found that lab-reared larvae have a depauperate gut community compared to wild individuals, suggesting that the larval gut *is* populated by microbes acquired from the environment (Xiang et al 2006; Pinto-Tomás and Sittenfeld 2011; Belda et al 2011). Although the adult butterfly gut probably is colonized at least in part by environmental microbes—as suggested by the high percentage of bacterial reads derived from OTUs also found in butterfly foods—the vast majority of ingested microbial species may fail to establish residence. The fact that the majority of the OTUs in butterfly foods were not detected in adult guts emphasizes the strength of the filter the gut exerts on potential microbial colonists, even in non-social hosts such as butterflies. Gut chemistry likely filters potential colonists: most of the microbes in the gut (excluding transients) will belong to the subset that can tolerate the gut’s pH and oxygenation conditions. Competition or priority effects from initial gut residents (those carried over from the larval host or established from early adult meals) could also prevent food-derived microbes from successful colonization. Both areas are fruitful avenues for future research aimed at understanding the community assembly of animal gut flora.

#### 3b. Known habitat requirements of dominant gut microbes suggest that the adult butterfly gut is acidic and potentially aerobic

The environmental requirements of the gut flora can help us infer chemical conditions in the adult butterfly gut, which can be difficult to measure using traditional means. Acetic acid bacteria (Acetobacteraceae) are obligately aerobic (Komagata et al 2014). The presence of many members of the Acetobacteraceae in the adult gut suggests that it is not entirely anoxic; oxygen may reach the outer layer of the gut via diffusion from gas in the tracheoles. Acetobacteraceae also prefer acidic conditions, as do the yeasts *Hanseniaspora*, *Candida* and *Kazachstania* (Rosa and Peter 2006; Komagata et al 2014). We could find no information about the pH of the adult butterfly’s gut, but in adults of the moth *Manduca sexta* it is between 5 and 6 (T. Hammer, unpublished data). The larval gut is often alkaline, ranging from close to neutral pH to over 12 (Gross et al 2008), but the prevalence of acetic acid bacteria and acid-tolerant yeasts in our data suggests that the adult gut is in fact acidic across most, if not all, butterfly species.

### 4. Functional potential of the butterfly gut flora

#### 4a. The catabolic potential of the gut flora varies with host diet

The catabolic capacity of the gut flora was related to host feeding ecology. The gut flora of frugivores exhibited increased catabolism of amino acids, carboxylic acids and dicarboxylic acids compared to nectivores’ gut flora, and decreased catabolism of sugars and sugar alcohols. These differences in function likely result from differences in the chemical composition of the diet. Nectars are 90% sugar by dry weight (Luttge 1977). Fruit generally contains more nitrogen than nectar: at our field site, the juice of rotting *Dipteryx oleifera*, the dominant food available to frugivorous butterflies during portions of the year, contains 33 times more essential amino acids and 19 times more non-essential amino acids per unit sugar than the flower nectars fed upon by butterflies (Ravenscraft and Boggs 2016). Frugivorous butterflies therefore provide a comparatively nitrogen-rich gut environment that appears to support microbes capable of catabolizing amino acids, while nectivores provide a more sugar-rich environment that favors microbes specialized on sugar catabolism.

A 96-well plate is a very different environment from a butterfly gut, therefore we cannot conclude that the functions we observed necessarily operate in vivo. However, our results indicate that host species and host diet are related not only to gut community composition, but also the functional capacities of the gut flora, and they suggest that the chemical makeup of the diet selects for microbes that are specialized to digest that diet.

### 5. Butterfly gut flora include potential mutualists, commensals and pathogens

A full understanding of the microbial gut community ultimately requires specific knowledge of each of its members. From what is known about the microbial genera present in the butterfly gut, we can begin to speculate whether they may be beneficial, commensal, or detrimental to butterfly hosts.

#### 5a. Bacteria

The genera *Enterobacter*, *Enterococcus*, and *Lactococcus* are commonly present in the digestive tracts of a wide range of animals, from mammals to insects, and likely participate in mutualistic or commensal relationships with their hosts (e.g. Robinson et al 2010; Engel and Moran 2013; Delsuc et al 2014). *Acetobacter, Asaia*, and *Pantoea* are frequent residents of insect guts (Crotti et al 2010; Engel and Moran 2013) and have been shown to directly benefit some hosts. For example, *Acetobacter pomorum* and *Asaia* promote dipteran larval development, probably through nutrient supplementation (Shin et al 2011; Mitraka et al 2013). *Pantoea agglomerans* helps the swarming locust, *Schistocerca gregaria*, synthesize aggregation pheromone (Dillon et al 2002). What these species may do for butterflies is currently unknown.

We also detected several genera that might be detrimental to butterflies. *Serratia* species are often insect pathogens (Grimont and Grimont 2006). The genus *Bartonella* contains opportunistic animal pathogens that can infect insects and mammals. Many are vectored by insects— primarily flies (Minnick and Anderson 2000)— but the genus has also been detected in the guts of other non-biting insects including honey bees, ants, and carrion beetles (Jeyaprakash et al 2003; Stoll et al 2007; Kaltenpoth and Steiger 2014). *Spiroplasma* and *Wolbachia* are well-known as reproductive parasites of insects; some kill males to promote their own spread via the female line. *Spiroplasma* are known to colonize the gut lumen. *Wolbachia* are generally associated with the reproductive organs, but have also been found in insect guts (Frost et al 2014; Berasategui et al 2016). We dissected out the guts of the butterflies in our study, but we were not always able to completely remove attached fat, Malphigian tubules, and reproductive tissue. Although we are confident that most of the OTUs reported here were gut-derived, the origin of the *Wolbachia* in our study is uncertain.

Although several of the most abundant bacterial genera we detected are often assumed to be harmful to their hosts and could well be detrimental to butterflies, evidence suggests these genera can sometimes be beneficial. *Spiroplasma* and *Wolbachia* can both protect their hosts from pathogens or parasites (e.g. Hedges et al 2008; Jaenike et al 2010) and *Wolbachia* has been shown to engage in nutritional symbioses, providing B vitamins to bedbugs (Hosokawa et al 2010). The high prevalence and vertical transmission of *Bartonella* in several fly species suggest that this genus could also be beneficial to some insect hosts (Halos et al 2004). Overall, butterflies host both potential mutualists and pathogens, but the balance seems shifted towards more beneficial than detrimental bacteria.

The data that do exist regarding the community composition of the adult butterfly gut flora are largely congruent with our findings. *Enterobacter*, *Enterococcus*, *Lactococcus, Commensalibacter* and *Orbus* were dominant members of the adult butterfly gut flora in both our data and in a study of the butterfly *Heliconius erato* in Panama (Hammer et al 2014). *Commensalibacter* has also been detected in *Drosophila* guts, where it may protect the flies from infection by pathogens (Roh et al 2008; Ryu et al 2008). It may play a beneficial role in butterflies, as well: in a related study, we found that abundance of *Commensalibacter* was positively correlated with increased adult lifespan in the temperate butterfly *Speyeria mormonia* (Ravenscraft et al, in review). Little is known about the biology of *Orbus*, but the genus or its close relatives have been detected in the guts of flies (Chandler et al 2011), bees (Kwong and Moran 2013), and geographically disparate butterflies including *Speyeria mormonia* in Colorado and *Sasakia charonda* in South Korea (Kim et al 2013). The apparently widespread associations between butterflies and bacteria in the genera *Commensalibacter* and *Orbus* suggest that these microbes may have a generalized relationship with lepidopterans or, more broadly, with sugar-feeding insects.

#### 5b. Fungi

Most of the fungal inhabitants of butterfly guts were saccharomycete yeasts. Members of the genus *Hanseniaspora* are early colonizers of decaying fruits and are also commonly isolated from drosophilid flies. They are dispersed by the flies, which also use them as a food source (Kurtzman et al 2011). The genera *Candida* and *Pichia* are globally widespread and present in many different environments. Some are pathogenic, but many are animal commensals or mutualists. *Candida* and *Pichia* species have been detected in the guts of both vertebrates and invertebrates including humans, gorillas, beetles, and larval lepidopterans (Rosa et al 1992; Suh et al 2005; Marchesi 2010; Dematheis et al 2012; Hamad et al 2014). All reported isolates of *Candida quercitrusa*, the species most closely related to the OTUs in our dataset, derive from flowers or insects (Kurtzman et al 2011). Not much is known about the ecology of *Kazachstania exigua*, but most strains have been isolated from human food products (Kurtzman et al 2011). While *Kazachstania* was present at artificially high abundance in our raw data due to its dominance in the trap baits, it was also detected on wild fruits and in the guts of butterflies that had not fed on trap baits, suggesting that it does naturally occur in butterfly guts at low abundance.

Many of the abundant fungal OTUs outside of the Saccharomycetales were too divergent from known species’ sequences to identify them with fine taxonomic resolution. One OTU was a member of the class Letiomycetes, a class of plant pathogens. Another belonged to the order Pleosporales, which predominantly consists of plant-decaying fungi, though some species associate with live plants (Zhang et al 2009). These fungi may have arrived in the adult gut via spores in fruits and nectars, or they could have colonized the larval gut via ingestion of leaf tissue and then persisted through pupation. In summary, the fungal flora of the butterfly gut included many sugar-loving yeasts and plant associates, some of which may pass through insect guts as accidental vagrants or hitchhikers, as well as potential insect gut-associated commensals or mutualists.

## Conclusions

Gut microbiota are a ubiquitous feature of animal life, yet we still understand little about the structure and function of these symbiotic communities. Here we developed adult Neotropical butterflies as a study system to partition variation in gut community membership at the levels of the host individual, species, and feeding guild; to compare the microbial species in the gut to those in host foods; and to investigate the catabolic potential of the gut flora. All samples were collected from a single site, thus eliminating the potential for large-scale geographic variation that is often present in, and can potentially confound, studies of similar scale.

We found that the majority of variation in the butterfly gut flora is expressed at the level of the individual host, followed by host species, with the least amount of variation explained by dietary guild. The consistency of these patterns across both bacterial and fungal communities lends support to the observed pattern. The pattern itself demonstrates the need for future work to expand from the current focus on broad diet categories and host taxonomy to additional, species-level or individual-level traits— for example, gut chemistry— in order to better understand the causes and consequences of the gut community structure in nature.

Butterfly gut flora had little resemblance to food microbial communities, suggesting that conditions in the gut exert a strong ecological filter on colonists. The initial colonists of the butterfly gut may be derived largely from butterfly foods, but once established, the butterfly gut community appears to be resistant to perturbation via ingestion of microbes. However, foods do influence the gut flora via their chemical composition: culture-based assays suggested that the chemical composition of the diet selects for gut microbes that are good at digesting common chemical components of that diet.

Butterfly guts host a rich community of bacteria, as well as yeasts, which are woefully understudied in contemporary analyses of the gut flora. By characterizing variation of the gut microbiota within and among butterfly species we have laid the foundation for a mechanistic understanding of how this hidden symbiosis affects and is affected by its host.

## Acknowledgments

Sincere thanks to Luke Frishkoff, Meredith Blackwell, and Tadashi Fukami’s 2013 Ecological Statistics class for insightful discussion and feedback. Jon Sanders provided the 16s rRNA standard for our qPCR standard curve. Carlos de la Rosa, Ronald Vargas, Bernal Matarrita Carranza, Danilo Brenes Madrigal and the staffs of OTS and La Selva Biological Station offered invaluable logistical assistance and support in the field. This work was supported by an NSF Graduate Research Fellowship, a Stanford Center for Computational, Evolutionary and Human Genomics Graduate Fellowship, and an NIH PERT postdoctoral fellowship to AR, funds from Stanford University, and grants from Stanford University’s Biology Department, Center for Latin American Studies, and Biosciences Office of Graduate Education.

## Data Accessibility

Raw Illumina sequences will be made available in the NCBI Short Read Archive. All other data (e.g. the OTU table, bacterial abundances calculated from qPCR, butterfly fecundity, egg chemical composition, etc) will be uploaded to Dryad.

